# HDL-associated phosphatidylserine blunts myeloid activation and protects from atherosclerosis

**DOI:** 10.1101/2025.11.26.689708

**Authors:** Aurora Bernal, Arslan Hamid, Cristina Grao-Roldán, Alice Scarpa, Iria Sánchez, José Ángel Nicolás-Ávila, Laura Pena-Couso, Mohamed H. Yaghmour, Alberto Benguría, Aránzazu Rosado, Lucía Sánchez-García, Carlos Torroja, Ana Dopazo, Fátima Sánchez-Cabo, Andrés Hidalgo, Max L. Senders, Mandy M.T. van Leent, Lea Seep, Jan Hasenauer, Katarzyna Placek, Christoph Thiele, Mihai G. Netea, Willem J.M. Mulder, Niels P. Riksen, Carlos Pérez-Medina

**Affiliations:** Centro Nacional de Investigaciones Cardiovasculares (CNIC), Madrid, Spain; Department for Immunology and Metabolism, Life and Medical Sciences Institute (LIMES), University of Bonn, Bonn, Germany; CIBER de Enfermedades Cardiovasculares, Madrid, Spain; Vascular Biology and Therapeutics Program and Department of Immunobiology, Yale University School of Medicine, New Haven, USA; BioMedical Engineering and Imaging Institute, Icahn School of Medicine at Mount Sinai, New York, New York, USA; Department of Surgery, Leiden University Medical Center, University of Leiden, Leiden, the Netherlands; Department of Diagnostic, Molecular, and Interventional Radiology, Icahn School of Medicine at Mount Sinai, New York, New York, USA; Department of Internal Medicine, Radboud University Medical Center, Nijmegen, The Netherlands; Department of Biomedical Engineering, Eindhoven University of Technology, Eindhoven, the Netherlands; Computational Life Sciences, Life and Medical Sciences Institute (LIMES), University of Bonn, Bonn, Germany

## Abstract

**Background and aims:** Lipids play a critical role in atherosclerosis. Low-density lipoprotein (LDL)-cholesterol and certain lipid classes like sphingomyelins are associated with inflammation and poor cardiovascular outcomes. Phosphatidylserine (PS), on the other hand, is a negatively charged anti-inflammatory phospholipid class involved in efferocytosis. In this study, we sought to investigate its anti-atherosclerotic properties through a combination of complementary human lipidomics analyses, *in vitro* assays and *in vivo* experiments in *Apoe^-/-^* mice.

**Methods:** Human lipidomics studies were performed on the 300OB cohort comprising 300 obese and overweight individuals at risk of cardiovascular disease. *In vitro* assays were carried out using human monocytes and macrophages, and *in vivo* experiments included histopathological, immunophenotyping and single-cell transcriptomic analyses.

**Results:** In humans, we identified PS as an anti-inflammatory and atheroprotective biomarker. Hence, we developed a high-density lipoprotein (HDL)-like formulation enriched in PS to exploit its properties in a targeted fashion in mice. *In vitro*, this formulation potently inhibited inflammatory cytokine production on human myeloid cells. Our in-depth *in vivo* experiments provided evidence of the formulation’s potent plaque-stabilizing and anti-inflammatory actions. These effects were mediated by a shift in the monocyte/macrophage compartment toward homeostatic/repairing phenotypes.

**Conclusions:** Collectively, our results demonstrate that HDL-associated PS potently suppresses inflammation and atheroprogression, and holds promise as a viable approach to improve immunomodulatory therapies.

## INTRODUCTION

Lipids play an essential role in atherosclerosis as key components of lipoproteins and plaques. Indeed, dyslipidemia, defined as increased circulating concentrations of triglycerides and cholesterol –especially when present as low-density lipoprotein (LDL)-cholesterol– is an established risk factor for cardiovascular complications. Likewise, sphingomyelins, their derivatives, and certain ceramides have also been associated with poor cardiovascular outcomes^1,2^. Lipoprotein accumulation and oxidation in the arterial wall drive inflammatory atherosclerosis^3^. The recruited pro-inflammatory monocytes differentiate into macrophages, which in turn become foam cells^4^ upon sustained uptake of oxidized lipoproteins. As this process continues over decades, early inflammatory lesions progress into fully-developed atherosclerotic plaques that may rupture and trigger atherothrombotic events, such as myocardial infarction (MI) or stroke^5^. While current pharmacological prevention or treatment of cardiovascular disease (CVD) mainly relies on lipid-lowering drugs, such as statins^4^, an increasing focus is currently devoted to inflammation^6,7^. Preclinical success in targeting inflammation in atherosclerosis models preceded the Canakinumab Anti-inflammatory Thrombosis Outcome Study (CANTOS), the first anti-inflammation trial in CVD patients. CANTOS showed that a treatment regimen involving an anti-interleukin 1β (IL-1β) antibody (canakinumab) reduced the incidence of recurrent cardiovascular events^8^. However, side effects in a number of patients, mostly related to severe infections, limit the applicability of this approach^9^. Recently, randomized trials using low-dose colchicine have also proved to decrease the incidence of cardiovascular complications^10–12^, further arguing for the importance of anti-inflammatory approaches in CVD.

At the crossroads of lipids and inflammation, high-density lipoprotein (HDL), a natural nanoparticle mainly composed of phospholipids, cholesterol and apolipoprotein A1 (apoA1), takes a central role^13^. HDL mediates reverse cholesterol transport^14^ and has additional anti-inflammatory and anti-oxidative properties^15^. Collectively, these features give HDL an intrinsic atheroprotective function, which consequently led to the development of HDL-raising interventions based on cholesteryl ester transfer protein (CETP) inhibitors, fibrates or HDL infusions^16^. However, these therapies did not improve standard-of-care treatment or had severe side effects^16,17^. Additionally, Mendelian randomization studies excluded a direct causal effect of natural HDL on CVD^18^. In this context, HDL’s phospholipid composition has been investigated as a possible variable in its behavior. For example, HDL enrichment with negatively charged phospholipids like phosphatidylserine (PS) correlated with higher cholesterol efflux capacity^19^ and enhanced anti-inflammatory effects on macrophages^19–21^. PS also has anti-inflammatory effects of its own as it is involved in the efferocytosis process as an “eat-me” signal^22^.

In this study, we sought to investigate the anti-atherosclerosis properties of PS. To that end, we first explored the association between circulating PS and atherosclerotic plaques in a cohort of 300 individuals with an increased risk for CVD due to obesity or being overweight. Indeed, lipidomics analysis identified PS as the strongest atheroprotective and anti-inflammatory factor among all lipid classes studied in this cohort. Subsequently, we devised a reconstituted HDL-like formulation incorporating PS, henceforth PS@rHDL, in order to harness its properties in a targeted fashion. After studying its physicochemical features and *in vitro* anti-inflammatory effects on human monocytes and macrophages, we tested PS@rHDL in a mouse model of atherosclerosis. Through an in-depth study based on histopathological analyses, immunophenotyping and single-cell transcriptomic profiling we provide solid evidence of its ability to inhibit inflammation and atheroprogression.

## METHODS

### SUBJECT DETAILS AND EXPERIMENTAL MODEL

#### Human participants

##### Cohort Overview

The 300OB cohort comprised 302 individuals between 55 and 82 years old with a body mass index (BMI) above 27 kg/m^2^ and has been described previously^23–26^. Recruitment was conducted at Radboud University Medical Center from 2014 to 2016. Lipid-lowering medication, when used, was interrupted 4 weeks prior to blood collection. None of the female participants was using hormone replacement therapy. Blood samples were collected in the morning after an overnight fasting. The study was approved by the Ethical Committee of the Radboud University (Nr. 46846.091.13). All participants gave written informed consent. Experiments were conducted according to the principles expressed in the Declaration of Helsinki.

##### Blood donors

Buffy coats from healthy donors were obtained from the University Hospital Bonn upon approval by the Ethik Commission (#009/21).

#### Mice

All animal care and procedures were based on an institutional protocol from the *Centro Nacional de Investigaciones Cardiovasculares* approved by the *Comunidad de Madrid (Consejería de Medio Ambiente y Ordenación del Territorio)*. Female *Apoe^−/−^* mice (B6.129P2-*Apoe^tm1Unc^*/J), acquired from Charles River Laboratories, were used in this study. Animals were kept at 20-24°C and 45-65% relative humidity. They were allowed food and water ad libitum. Eight-week-old *Apoe^−/−^* mice were fed a Western diet (WD; high-cholesterol, high-fat diet: 0.21% weight cholesterol; 15.2% kcal protein, 42.7% kcal carbohydrate, 42.0% kcal fat; Ssniff, E15721-347 EF TD88137) for 6 (WD_6w_) or 14 (WD_14w_) weeks to obtain atherosclerotic lesions at differing disease stages. These mice present high LDL and total cholesterol concentrations in blood resulting from their lack of apolipoprotein E, leading to the accelerated development of atherosclerotic lesions^27,28^.

In WD_6w_ mice, we consistently found established fibroatheroma lesions in the aortic valve, and smaller plaques were starting to develop in the arch and bifurcation. The descending aorta, more atheroresistant, was lesion-free at this stage. WD_14w_ mice, on the other hand, showed more advanced plaques in the aortic root with calcifications while lesions in the arch and bifurcation showed an advanced phenotype similar to WD_6w_ valve lesions. In WD_14w_ we observed incipient, early-stage lesions in the descending aorta. Age-matched female C57BL/6J mice fed a normal chow diet for an equal period of time were used as healthy wild type (WT) controls. Male DsRed (CD45.2) and male BL/6.SJLC57 (CD45.1) mice were used for *in vivo* phagocytosis analysis. A minimum of 10 animals (n = 10) per group was considered in every experiment, unless otherwise specified.

### QUANTIFICATION AND STATISTICAL ANALYSIS

Data were analyzed for normality using the Kolmogorov-Smirnov test. Normally distributed data were analyzed using unpaired two-tailed student’s t test for comparisons between two groups, or one-way analysis of variance (ANOVA) to evaluate differences between the PS@rHDL group and the other groups, followed by multiple comparisons using Dunnett’s test. These data are expressed as mean ± standard deviation (SD). Non-parametric Kruskal-Wallis analysis followed by multiple comparisons using Dunn’s test was performed to assess differences between the PS@rHDL group and the other groups on the data that did not pass the normality test. These data are presented as median with interquartile range (IQR). Fisher’s exact test was used for qualitative analysis of calcifications (Alizarin red staining). Gehan-Breslow-Wilcoxon test was performed for survival analysis. Statistical analyses were performed with GraphPad Prism software version 9.4.1 and differences with P values < 0.05 were considered significant, shown in results as * P < 0.05, ** P < 0.01, ***, P < 0.001 and **** P < 0.0001.

### METHOD DETAILS

Method details are described in the **Supplementary material**, including **human lipidomics** (cardiovascular measurements, metabolic parameters, measurement of circulating lipids, peripheral blood cell isolation and stimulation), **lipidomic analysis** (chemical solvent and reagent quality assurance, sample preparation, instrumental configuration, data acquisition and data processing and analysis), **human *in vitro* procedures** (human PBMC and monocyte isolation, PBMCs and monocytes in vitro procedures and cytokine quantification), **nanoformulation** (chemicals, synthesis and characterization of apoA1 formulations and radiolabeling of PS@rHDL with ^89^Zr), ***in vitro* murine procedures** (cholesterol efflux assay and efferocytosis *in vitro*), **animal procedures** (pharmacokinetics, biodistribution, targeting assay, treatment schedule, toxicity, ^18^F-FDG study, in vivo phagocytic activity quantification and infection challenge experiment), ***ex vivo* procedures** (flow cytometry, histology and immunohistochemistry, single-cell RNA sequencing (scRNA-seq) and scRNA-seq bioinformatics analysis).

## RESULTS

### Plasma PS concentrations are negatively associated with atherosclerosis and inflammation in obese individuals

Using direct infusion-mass spectrometry (DIMS) we analyzed the plasma lipid composition in a cohort of 300 obese and overweight individuals (300OB cohort), that has been described before^23–26^. Fifty-three percent of these individuals had carotid atherosclerotic plaques (Fig. 1A). Among the 17 identified lipid classes (Table S1), triglycerides, diglycerides, sphingomyelins and ceramides showed strong positive associations with intima media thickness (IMT) values, maximal plaque thickness and number of plaques, consistent with their detrimental effect on CVD^1,2^ (Fig. 1B). Notably, PS concentrations showed the strongest significant negative correlations with carotid IMT, maximal thickness and number of plaques, suggesting a potential protective role against plaque formation. PS concentrations showed a strong positive correlation with HDL-cholesterol (Fig. 1C), which may reflect an enrichment of this phospholipid in certain HDL fractions^19^. Interestingly, we observed a strong negative correlation between PS and a panel of circulating inflammatory markers including IL-18, IL-18 binding protein (IL-18bp), resistin and alpha-1-antitrypsin (AAT) (Fig. 1D). Furthermore, PS plasma concentrations negatively correlated with cytokine IL-1β, TNF-α and IL-6 production capacity by peripheral blood mononuclear cells (PBMCs) upon stimulation with Pam3Cys and lipopolysaccharide (LPS) (Fig. 1E), although the associations did not reach statistical significance. These findings suggest potent inhibitory effects of PS on inflammatory processes in obese and overweight individuals.

**Figure 1.**
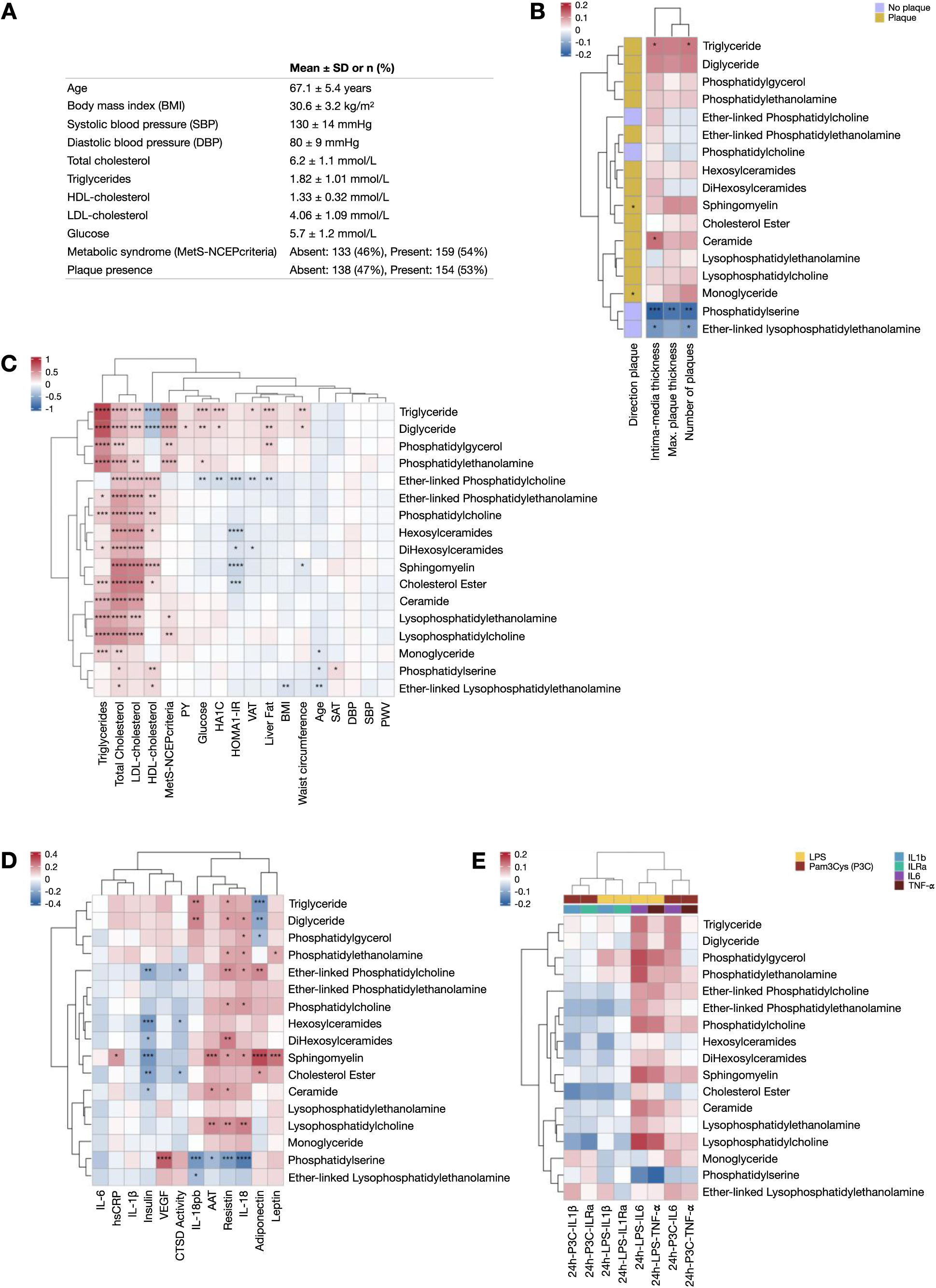
Plasma PS concentrations are negatively associated with atherosclerosis and inflammation in obese individuals. **A**. Baseline characteristics of the 300OB Cohort. **B-E.** Correlations between 17 lipid classes and clinical and inflammatory parameters of the patients, including carotid atherosclerosis determined by plaque presence, carotid intima-media thickness (IMT), maximum plaque thickness, and total plaque count (**B**); other clinical and risk factors (**C**); circulating inflammatory proteins and adipocytokines (**D**); and cytokine production capacity by PBMCs stimulated with lipopolysaccharide (LPS) or Pam3Cys-Ser-(Lys)4 (P3C) for 24 hours, specifically interleukin 1β (IL-1β), interleukin 1 receptor antagonist (IL-1Ra), interleukin 6 (IL-6), and tumor necrosis factor-alpha (TNF-α) (**E**). All correlations were determined using Spearman’s rank correlation coefficient, except for correlations of lipid class concentrations and the presence of atherosclerotic plaques where the Wilcoxon Rank Sum test was used. * P < 0.05, ** P < 0.01, *** P < 0.001, and **** P < 0.0001. PY: number of pack years; VAT: visceral adipose tissue volume; SAT: subcutaneous adipose tissue volume; DBP: diastolic blood pressure; SBP: systolic blood pressure; PWV: pulse wave velocity. PBMCs: peripheral blood mononuclear cells.

### HDL-associated PS potently inhibits inflammatory cytokine production in human monocytes and macrophages

Spurred by these results, we designed a reconstituted HDL-like formulation enriched in PS (PS@rHDL, Fig. 2A), in order to exploit the observed features of PS in a targeted fashion. As control nanoformulation, we also synthesized the PS-free counterpart (rHDL, Fig. 2A). The two formulations had similar chromatographic retention times (Fig. 2B) and mean hydrodynamic sizes of approximately 10 nm (Fig. S1A). We conducted *in vitro* studies to assess PS@rHDL’s anti-inflammatory action on human PBMCs (Fig. 2C) and macrophages differentiated in the absence (Fig. 2D) and presence of oxidized LDL (oxLDL, Fig. 2E). In these assays we measured IL-6 and TNF-α production in response to Pam3Cys and LPS, and observed some anti-inflammatory effects for PS and rHDL. However, while both formulations had different effects on cytokine production depending on stimulus and cell type, PS@rHDL consistently showed a potent inhibitory action on IL-6 and TNF-α production in all tested conditions. We further explored PS@rHDL’s properties *in vitro* using murine bone marrow-derived macrophages (BMDMs). PS@rHDL efficiently increased cholesterol efflux (Fig. S1B) and enhanced efferocytosis (Fig. S1C) compared to PS and rHDL. Taken together, these results suggest a synergistic effect resulting from PS incorporation into an HDL-like formulation.

**Figure 2.**
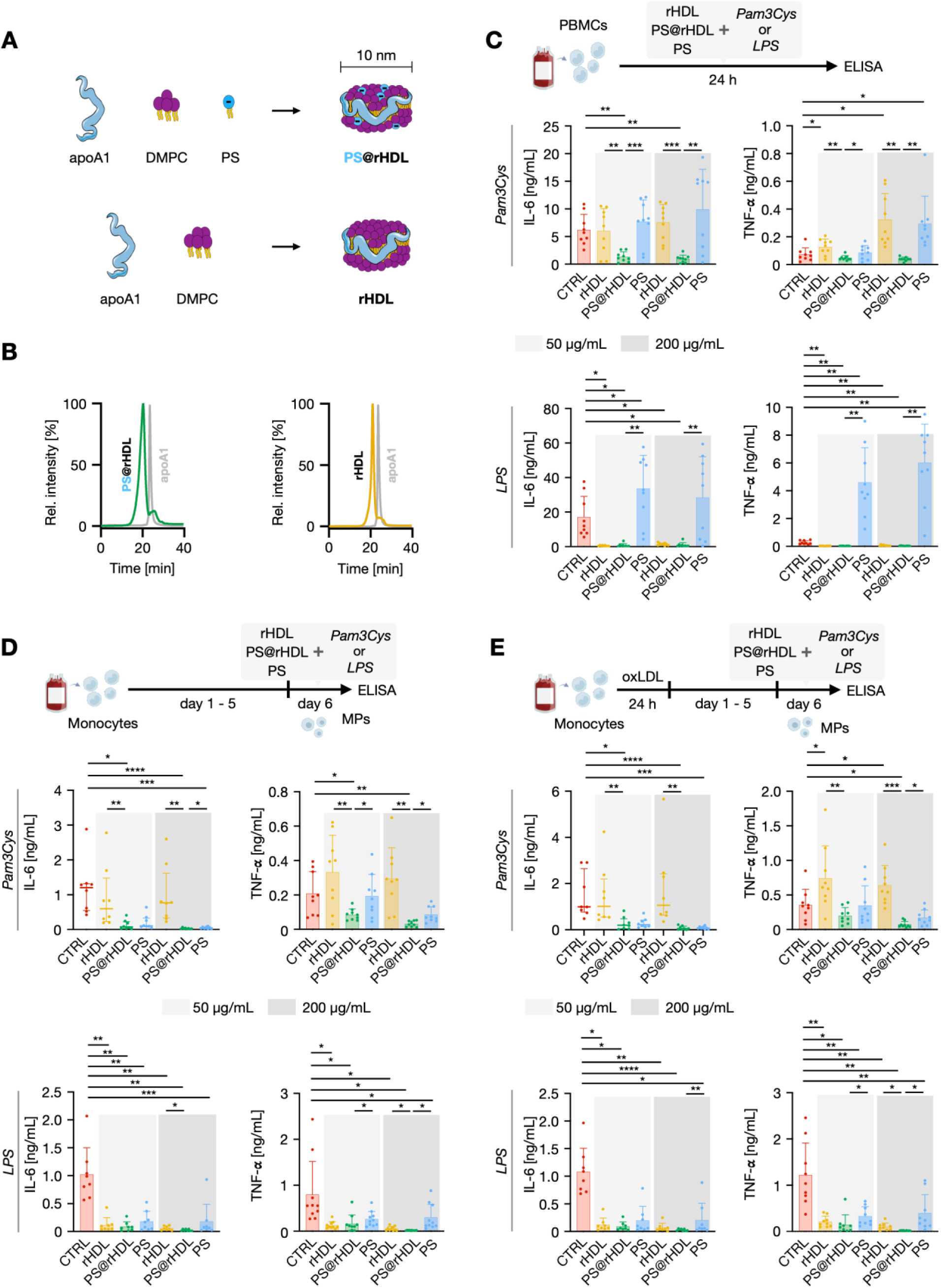
HDL-associated PS potently inhibits inflammatory cytokine production in human monocytes and macrophages. **A.** Schematic of PS@rHDL and rHDL formulations. **B**. Size exclusion chromatograms showing PS@rHDL (green) and rHDL (yellow). The apoA1 chromatogram (grey) is shown for reference. **C.** IL-6 and TNF-α production by PBMCs treated with PBS, rHDL, PS@rHDL or PS, after Pam3Cys or LPS stimulation **D.** IL-6 and TNF-α production by macrophages 6 days post differentiation treated with PBS, rHDL, PS@rHDL or PS and stimulated with Pam3Cys or LPS. **E.** IL-6 and TNF-α production by macrophages 6 days post differentiation, after an initial 24-hour exposure to oxLDL, treated with PBS, rHDL, PS@rHDL or PS and re-stimulated with Pam3Cys or LPS. Normally distributed data are presented as mean ± SD, and P values were calculated using one-way ANOVA, followed by multiple comparisons using Dunnett’s test. Otherwise, data are presented as median and IQR, with P values calculated using Kruskal-Wallis test, followed by multiple comparison using Dunn’s test. * P < 0.05, ** P < 0.01, *** P < 0.001 and **** P < 0.0001. DMPC: 1,2-dimyristoyl-*sn*-glycero-3-phosphocholine; PBMC: peripheral blood mononucleated cells; LPS: lipopolysaccharides; MPs: macrophages.

### PS@rHDL reduces necrotic core and plaque size in atherosclerosis

For *in vivo* studies, we used *Apoe^-/-^* mice fed a high-fat, high-cholesterol diet (Western diet, WD) for either 6 (WD_6w_) or 14 weeks (WD_14w_, Fig. 3A, left). Using Zirconium-89 (^89^Zr)-labeled PS@rHDL, we found high bone marrow, liver, and spleen uptake in both groups (Fig. S1D), in agreement with our previous work using similar formulations^29^. We then investigated PS@rHDL’s specificity for immune cell populations in aorta, blood, bone marrow and spleen single-cell suspensions by flow cytometry using DiR-labeled PS@rHDL (Fig. S1E). In all tissues, PS@rHDL showed a marked selectivity for myeloid cells (Fig. S1F). Additional *in vivo* targeting features are reported in the Supplemental Information section (Fig. S1G-H, Table S2). Next, we evaluated the anti-atherosclerotic effects of PS@rHDL on plaques at different disease stages, with special focus on monocyte/macrophage (Mo/MP) infiltration in the earlier stage (WD_6w_) and on the evolution of calcifications in the later stage (WD_14w_). The treatment regimen consisted of 8 administrations over two weeks (Fig. 3A, right) and was well tolerated (Fig. S2 & S3). Importantly, no differences were observed in circulating cholesterol concentrations in either mouse cohort (Fig. S2D & S3D), indicating that the observed effects are not due to changes in the exposure to cholesterol. Histological analysis of aortic valves from WD_6w_ mice treated with PS@rHDL showed reduced plaque size compared to controls, which was mainly due to a significant reduction in necrotic core area (Fig. 3B, S4A-B). These changes resulted in a larger tunica media thickness for PS@rHDL-treated mice compared to controls (Fig. 3B, S4C). We found, however, no apparent change in collagen content (Fig. S4D-E). Staining with antibodies against Mac-2 as a marker for inflammatory macrophages and foam cells, showed reduced area in plaques from WD_6w_ mice treated with PS@rHDL (Fig. 3C, S4F). No sign of calcification was found in plaques from WD_6w_ mice (Fig. S4G). The WD_14w_ groups overall showed larger plaques in the aortic root (Fig. 3D) compared to WD_6w_ mice. PS@rHDL treatment led to mild reductions in plaque size (significantly when compared to PS; Fig. 3D, S4H), and a marked reduction in necrotic core area (Fig. 3D, S4I). We found no differences in media thickness (Fig. 3D, S4J). As in WD_6w_ mice, plaque collagen content was similar in all groups (Fig. S4K, L), whereas no differences were observed in Mac-2^+^ area at this stage (Fig. S4M). We did observe micro– and macrocalcifications in a number of samples from saline (PBS)-, PS-, and rHDL-treated mice, but not in mice treated with PS@rHDL (Fig. 3E). Elsewhere in the aortic arch, both WD_6w_ and WD_14w_ mice showed smaller and less advanced lesions compared to the plaques found in the aortic root (Fig. S5). A similar trend toward reduced lesion size in PS@rHDL-treated animals in both WD_6w_ and WD_14w_ mice was found, although in most cases the differences observed did not attain statistical significance due to the greater variability found in these regions (Fig. S5).

**Figure 3.**
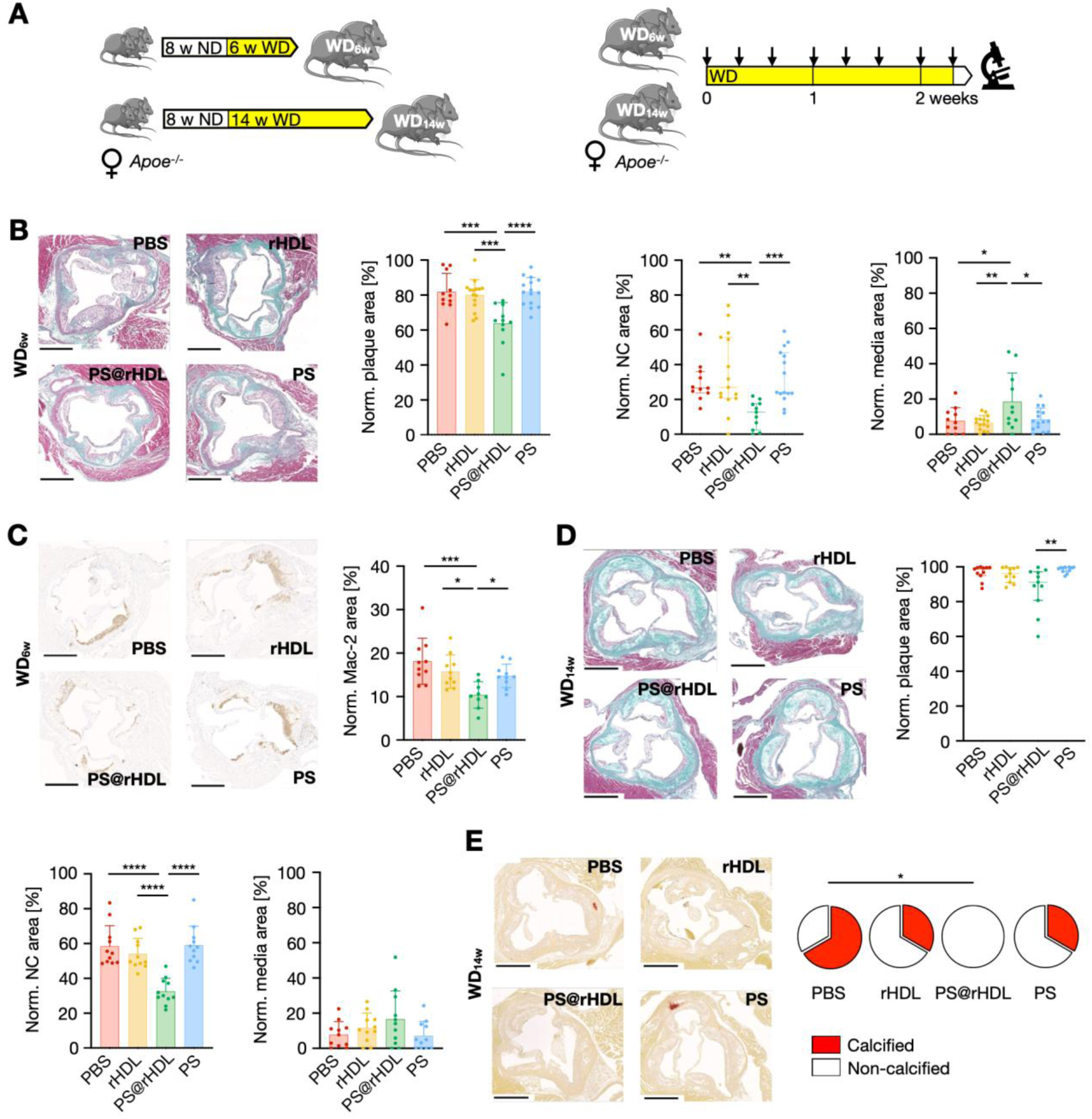
PS@rHDL reduces necrotic core and plaque size in atherosclerosis. **A.** The *Apoe^-/-^* mouse atherosclerosis model was used in this study. Female *Apoe^-/-^* mice were fed a high-fat, high-cholesterol diet (Western diet, WD) for 6 (WD_6w_) or 14 weeks (WD_14w_) to attain atherosclerotic lesions of differing degree of disease progression (left). WD_6w_ or WD_14w_ mice were treated (right) with PBS (saline control), rHDL, PS@rHDL or PS over 16 days receiving 8 treatment injections (arrows). Animals were euthanized 24 hours after the last administration for sample collection and analysis. **B.** Representative images of atherosclerotic lesions in the aortic root from treated WD_6w_ mice stained with Masson’s Trichrome. Scale bar = 500 µm. Quantitative analysis of normalized plaque, normalized NC and normalized tunica media areas in the aortic root from treated WD_6w_ mice (n = 10-15 per group). **C.** Representative immunohistochemistry images (left) and quantitative analysis (right) of normalized Mac-2^+^ area of atherosclerotic plaques in the aortic root from treated WD_6w_ mice (n = 9-10 per group). Scale bar = 500 µm. **D.** Representative images of atherosclerotic plaques in the aortic root from treated WD_14w_ mice stained with Masson’s Trichrome. Scale bar = 500 µm. Quantitative analysis of normalized plaque, normalized NC and normalized tunica media areas in the aortic root from treated WD_14w_ mice (n = 11 per group). **E.** Representative images of Alizarin red-stained sections from atherosclerotic lesions in the aortic root from treated WD_14w_ mice (n = 8-9 per group). Scale bar = 500 µm. Additionally, pie charts are displayed indicating the share of samples showing calcifications (red area). Normally distributed data are presented as mean ± SD, and P values were calculated using one-way ANOVA, followed by multiple comparisons using Dunnett’s test. Otherwise, data are presented as median and IQR, with P values calculated using Kruskal-Wallis test, followed by multiple comparison using Dunn’s test. Fisher’s exact test was used for qualitative analysis of calcification (Alizarin red staining). * P < 0.05, ** P < 0.01, *** P < 0.001 and **** P < 0.0001. WD_6w_: *Apoe*^−/−^ mice fed a WD for 6 weeks; WD_14w_: *Apoe*^−/−^ mice fed a WD for 14 weeks; NC: necrotic core.

### PS@rHDL normalizes vessel wall metabolism and architecture

We investigated the effects of PS@rHDL treatment in the aorta at the tissue level. A significant reduction in aortic mass was measured in PS@rHDL-treated animals versus controls in both WD_6w_ and WD_14w_ mice independent of body mass (Fig. 4A & S6A), closely approaching values from healthy wild type (WT) mice (indicated by dashed lines in Fig. 4 graphs). Additionally, we investigated metabolic adaptations after PS@rHDL treatment in the vessel wall using [^18^F]-2-deoxy-2-fluoro-D-glucose (^18^F-FDG). A significantly lower ^18^F-FDG uptake was measured in aortas from WD_6w_ mice treated with PS@rHDL as compared with saline and PS controls, although the differences with rHDL-treated animals were not significant (Fig. 4B left, S6B). In WD_14w_ mice there was a trend towards reduced uptake in PS@rHDL-treated mice compared to saline controls (P = 0.10, Fig. 4B right), and the observed differences were significant for uptake values that were dose-corrected only (%ID, Fig. S6C). In both cohorts, the uptake values of the PS@rHDL groups approached those of healthy WT mice. We next performed autoradiography on whole aortas to assess regional ^18^F-FDG uptake distribution (Fig. S6D). Analysis of autoradiographs showed that aortic arch and root (AAR)-to-descending aorta (DA) uptake ratios from PS@rHDL-treated mice approached healthy WT values in both cohorts (Fig. S6E). Taken together, these data reveal a decrease in tissue glycolytic metabolism in aortas from mice treated with PS@rHDL, both in the plaque-rich AAR and the plaque-free DA wall. No major differences were found in other relevant tissues (Fig. S6F-J). We subsequently reasoned that these observations might be due to a systemic effect of the treatment on the whole arterial wall and not only at lesion sites, and decided to investigate histological changes. We thus measured different parameters of arterial wall architecture in the lesion-free descending thoracic aorta at the two points indicated in Fig. 4C, using three different stains (Fig. 4D). WD_6w_ mice treated with PS@rHDL had reduced vessel wall thickness, collagen burden and number of elastin fiber breaks, and this was accompanied by an increase in the number and density of nuclei in the upper descending thoracic aorta (Fig. 4D). Remarkably, the resulting values for all the measured parameters after PS@rHDL treatment were close to those of healthy WT controls. Similar results were observed in WD_14w_ mice (Fig. 4E), and in the lower descending thoracic aorta of both cohorts (Fig. S7). Collectively, these results suggest potent protective properties of PS@rHDL on arterial wall integrity.

**Figure 4.**
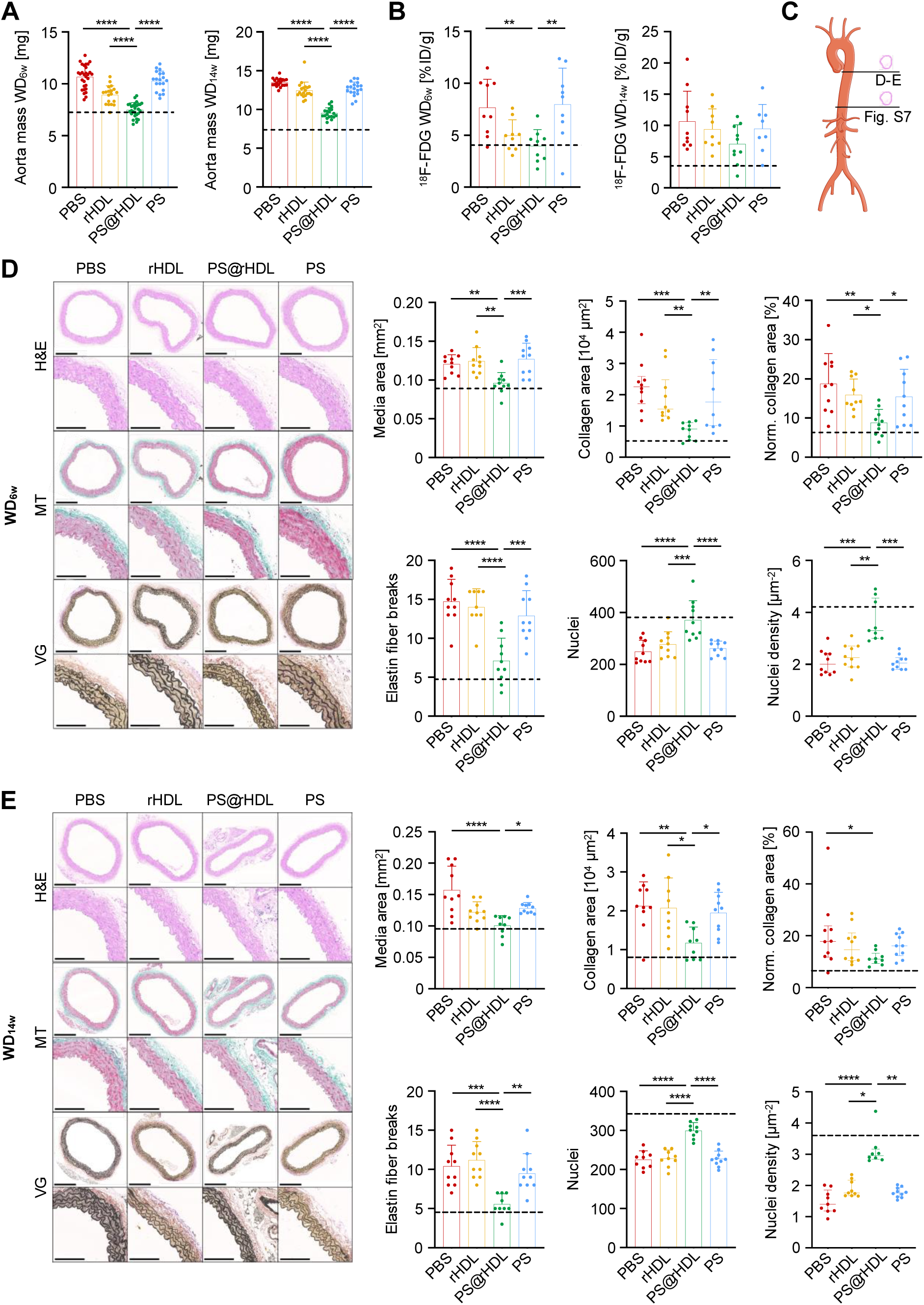
PS@rHDL normalizes vessel wall metabolism and architecture. **A.** Aorta mass after treatment for WD_6w_ (left, n = 20-30 per group) and WD_14w_ mice (right, n = 20 per group). **B.** ^18^F-FDG aortic uptake in the aortas for WD_6w_ (left, n = 9 per group) and WD_14w_ mice (right, n = 9 per group) after treatment. **C.** Schematic representation of the upper and lower descending thoracic aorta sections used for vessel wall histological analysis. **D.** Representative images showing hematoxylin & eosin (H&E), Masson’s Trichrome (MT) and Van Gieson (VG) stainings of the vessel wall from the upper section of the descending thoracic aorta from treated WD_6w_ mice (scale bar = 250 μm) and their respective magnifications (scale bar = 100 μm) after treatment. Quantitative analysis of tunica media area, collagen area, normalized collagen area, number of elastin fiber breaks, number of nuclei and nuclei density in the upper section of the descending thoracic aorta from treated WD_6w_ mice (n = 10 per group). **E.** Representative images showing hematoxylin & eosin (H&E), Masson’s Trichrome (MT) and Van Gieson (VG) stainings of the vessel wall from the upper section of the descending aorta from treated WD_14w_ mice (scale bar = 250 μm) and their respective magnifications (scale bar = 100 μm). Quantitative analysis of tunica media area, collagen area, normalized collagen area, number of elastin fiber breaks, number of nuclei and nuclei density in the upper section of the descending thoracic aorta from treated WD_14w_ mice (n = 10 per group). Dashed black lines in graphs represent average values from age-matched healthy wild type C57BL/6J mice. Normally distributed data are presented as mean ± SD, and P values were calculated using one-way ANOVA, followed by multiple comparisons using Dunnett’s test. Otherwise, data are presented as median and IQR, with P values calculated using Kruskal-Wallis test, followed by multiple comparison using Dunn’s test. * P < 0.05, ** P < 0.01, *** P < 0.001 and **** P < 0.0001. WD_6w_: *Apoe^−/−^* mice fed a WD for 6 weeks; WD_14w_: *Apoe^−/−^* mice fed a WD for 14 weeks; H&E: hematoxylin & eosin; MT: Masson’s Trichrome; VG: Van Gieson.

### PS@rHDL modulates the monocyte/macrophage compartment

In an initial analysis of Mo/MP populations by flow cytometry (Fig. S8A), we used two canonical markers of inflammatory status on myeloid cells, namely Ly6C and CD206, in order to assess potential phenotype changes due to PS@rHDL treatment. In the WD_6w_ aortas we did not observe differences in lineage positive cell (which included CD90.2^+^ and CD49b^+^ T cells, NK1.1^+^ natural killer cells, B220^+^ B cells and Ter119^+^ erythrocytes, Fig. S8B) or myeloid cell (Fig. S8C) frequencies among the different treatment groups. However, while there was no difference in neutrophil (Fig. S8D), dendritic cell (Fig. S8E) or monocyte frequencies (Fig. S8F), there was a marked decrease in Ly6C^hi^ monocyte percentages in PS@rHDL-treated animals (Fig. S8G). Similarly, we found no differences in macrophage burden (Fig. S8H), measured either as number of cells or frequency, but we did find an increase in CD206^+^ cells within this compartment (Fig. S8I). In WD_14w_ aortas we observed an overall decrease in lineage-positive cell frequencies (Fig. S9A) with a concomitant increase in myeloid cell percentages (Fig. S9B) compared to WD_6w_ mice. Likewise, while we found no changes in neutrophil and dendritic cell percentages (Fig. S9C-D), PS@rHDL induced the same modulatory effect within the monocyte (Fig. S9E-F) and macrophage (Fig. S9G-H) compartments, *i.e.*, a decrease in Ly6C^hi^ and an increase in CD206^+^ cells, respectively. The same pattern was observed in bone marrow (Fig. S10), blood (Fig. S11), and spleen (Fig. S12) from both groups after PS@rHDL treatment, with a decrease in Ly6C^hi^ monocyte and an increase in CD206^+^ macrophage percentages, suggesting that PS@rHDL globally antagonized the inflammatory profile of the myeloid cell compartment.

### scRNA-seq maps vessel wall and plaque cellular heterogeneity

Having clearly established the differences between PS@rHDL and all control treatments regarding their anti-atherosclerosis properties, we next sought to gain a deeper insight into the effect of incorporating PS into an HDL-like formulation using a simplified experimental paradigm comparing PS@rHDL and rHDL. To that end, we performed scRNA-seq on atherosclerotic aortic root and arch samples from WD_6w_ mice after treatment with these two formulations. Unbiased clustering yielded 16 cell populations (Fig. S13A), including smooth muscle cell (SMC, 3 clusters: C0, C4, C9), endothelial cell (EC: C8, C10, C11), immune cell (IC: C5, C12, C14), fibroblast (FB: C1, C2, C3, C6, C7, two of them –C3, C6– previously identified as valvular interstitial cells^27^), mesothelial cell (C13) and neuron (C15) populations. This clustering is in concordance with recent scRNA-seq studies in *Apoe^-/-^* mice^30–33^. A selection of differentially expressed genes for each cluster is shown in Fig. S13B (see Fig. S14A for full dot plot). Within the SMC compartment, an *Fn1*– and *Lgals3*-expressing subpopulation (C4) showing decreased expression of canonical contractile genes (*Acta2*, *Myh11*, *Myl9*) was identified and corresponds with a transitional, modulated population described by Wirka *et al*.^32^ (Fig. S13C). We also found a cluster (C2) expressing stem cell (*Ly6a*), endothelial cell (*Vcam1*) and monocyte (*Ly6c1*) genes, hence termed SEM cells in previous studies^33,34^. These cells also express *Il1r1*, *Csf1* and ECM remodeling (*Dcn*, *Lum*) and collagen (*Col1a1*) genes, which suggests they are an SMC/FB transitional cell state (Fig. S13C). An EC subpopulation (C8) highly expressing genes related to lipid handling (*Cd36*, *Scarb1*, *Fabp4*) was also identified. This EC subpopulation expands in response to WD^30,31^. Within the IC clusters, we found a large Mo/MP subpopulation (C5) along with smaller dendritic cell (DC, C12) and T cell (C14) clusters. Re-clustering of the myeloid populations (C5 and C12) resulted in 3 Mo/MP (C0_M_, C1_M_, C2_M_) and 1 DC cluster (C3_M_) (Fig. S13D, see Fig. S14B for full dot plot). Within the Mo/MP subpopulations, C0_M_ expressed genes typically associated with inflammatory Mo/MPs (*Il1b*, *Ccr2*, *Lgals3*); C1_M_ was enriched in genes expressed by resident-like/alternatively activated macrophages (*Mrc1*, *Cd163*, *Lyve1*, *Folr2*); and C2_M_ contained proliferating Mo/MPs (Fig. S13E). Importantly, most inflammatory and proliferating Mo/MPs expressed *Cx3cr1*, suggesting their previous recruitment from circulating blood^35,36^ (Fig. S13F).

### scRNA-seq reveals population changes between PS@rHDL– and rHDL-treated samples

Analysis of the scRNA-seq data by treatment condition showed differences in relative subpopulation weights within the identified cell types (Fig. 5A). Overall, there were only small changes among the four main cell populations (Fig. 5B). Functional enrichment analysis revealed an overall downregulation in collagen and extracellular matrix (ECM) organization genes, as well as inflammatory response genes in the PS@rHDL group (Fig. 5C). Further analysis of population changes within cell types revealed differences by treatment condition (Fig. 5D). Within SMCs, a relative reduction in the modulated *Lgals3*-expressing subpopulation (C4, Fig. S14C) was observed in the PS@rHDL group, with a concomitant increase in the contractile cluster (C0). Similarly, in the FB compartment, C2’s (SEM cells) relative size decreased after PS@rHDL treatment, with accompanying reduction in the expression of *Il1r1*, *Lgals3*, *Lum* or *Mmp3* genes. Within the EC subpopulations we observed a reduction in WD-associated C8’s relative size, while C10 and C11 both expanded after PS@rHDL treatment. Finally, a relative expansion of the Mo/MP cluster (C5) was observed at the expense of DCs (C12). Functional enrichment analysis in C2 (SEM cells) revealed a downregulation of genes involved in ECM organization, monocyte chemotaxis or IL-1 response (Fig. 5E) in the PS@rHDL group, whereas in C4 (modulated SMCs) we found a downregulation of integrin activation and cellular response to LDL genes, and upregulation of muscle contraction-related genes (Fig. 5F). Among myeloid cells, a rebalancing between inflammatory Mo/MPs and resident-like/alternatively activated MPs was observed (Fig. 6A & B), in line with our flow cytometry results. Furthermore, beside the abovementioned reduction in the relative size of the DC cluster, we found a reduced proliferating Mo/MP subpopulation in the PS@rHDL group (C2_M_, Fig. 6B). These population changes were paralleled by a decrease in inflammatory gene expression and a consequent increase in the expression of genes associated with resident-like/alternatively activated macrophages (Fig. 6C), typically associated with homeostatic phagocytosis. Functional enrichment analysis in these cell clusters show an overall downregulation of inflammatory pathways (NF-kB, prostaglandins) and an upregulation of genes related to receptor-mediated endocytosis and lipoxin synthesis, collectively suggestive of inflammation resolution (Fig. 6D).

**Figure 5.**
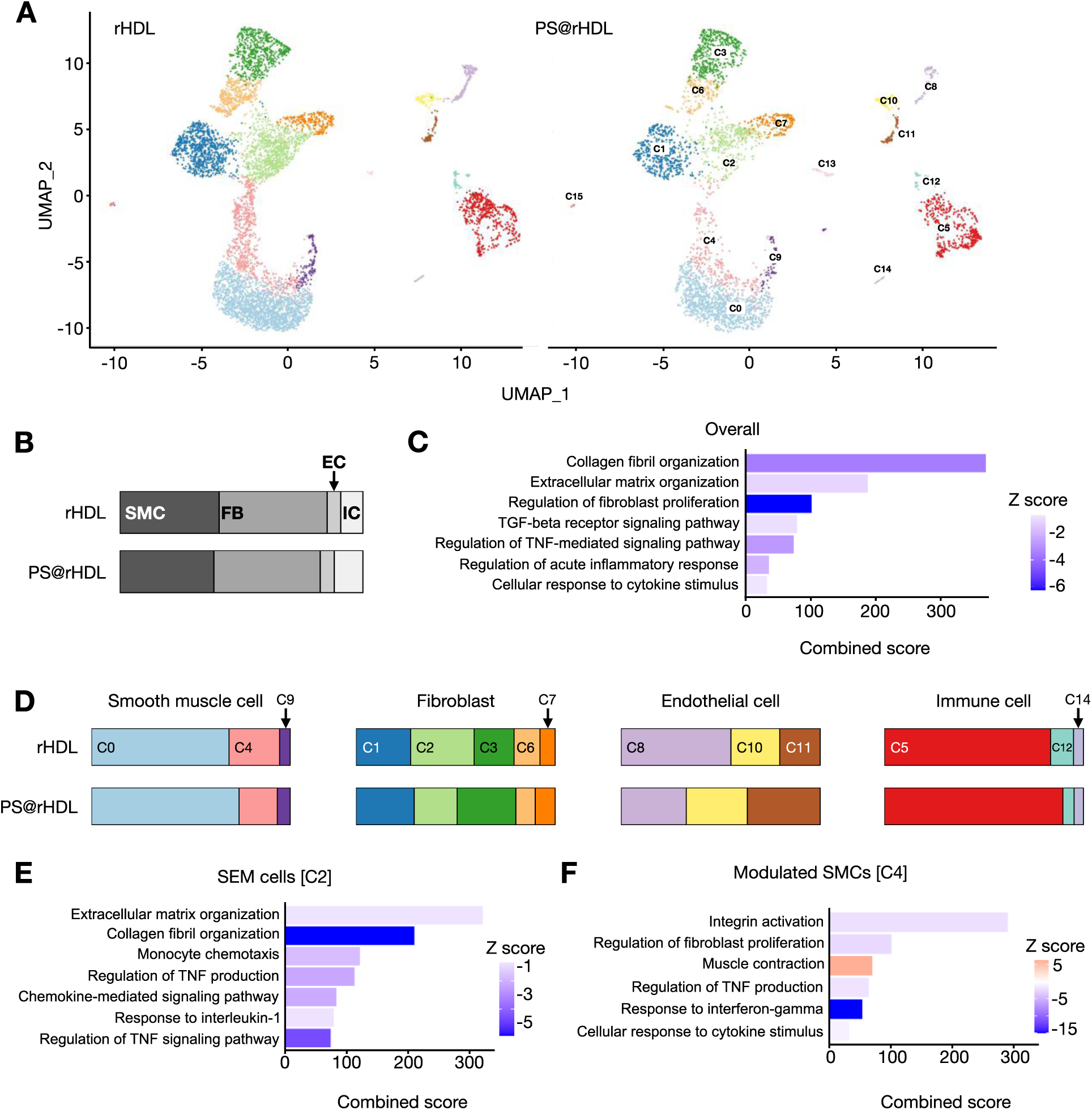
scRNA-seq reveals population changes between the PS@rHDL– and rHDL-treated groups. **A.** UMAP plots of single cells from aortic root and arch samples from WD_6w_ mice treated with rHDL (left) and PS@rHDL (right). **B.** Relative proportion of major cell types, *i.e.*, smooth muscle cells (SMC), fibroblasts (FB), endothelial cells (EC), and immune cells (IC), in each treatment group. **C.** Selected gene ontology (GO) terms with P_adj_ < 0.05 enriched in an overall analysis based on treatment condition (rHDL vs. PS@rHDL) including all clusters. Blue indicates downregulation in PS@rHDL vs. rHDL, red indicates upregulation in PS@rHDL vs. rHDL. **D.** Relative proportion of clusters within major cell type populations, in each treatment group. **E, F.** Selected GO terms with P_adj_ < 0.05 enriched in SEM cells (C2) (**E**) and modulated SMCs (C4) (**F**). Blue indicates downregulation in PS@rHDL vs. rHDL, red indicates upregulation in PS@rHDL vs. rHDL. WD_6w_: *Apoe^−/−^* mice fed a WD for 6 weeks.

**Figure 6.**
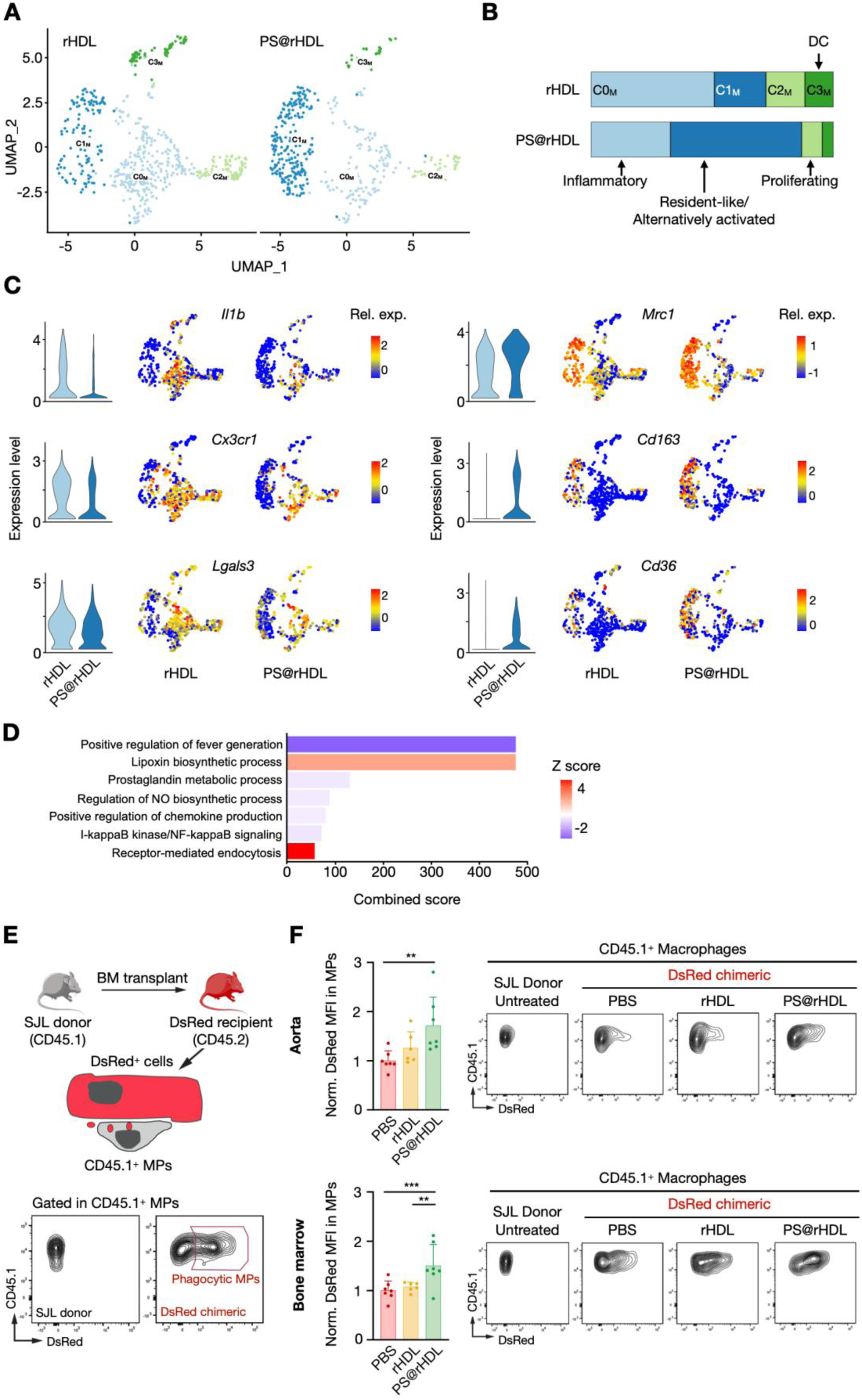
scRNA-seq reveals a myeloid switch between the PS@rHDL– and rHDL-treated groups. **A.** UMAP plots of single myeloid cells from aortic root and arch samples from WD_6w_ mice treated with rHDL (left) and PS@rHDL (right). **B.** Relative proportion of clusters in each treatment group. **C.** Violin and UMAP plots for selected representative genes showing relative expression according to treatment condition. **D.** Selected GO terms with P_adj_ < 0.05 enriched in an overall analysis based on treatment condition (rHDL vs. PS@rHDL) including all myeloid clusters. Blue indicates downregulation in PS@rHDL vs. rHDL, red indicates upregulation in PS@rHDL vs. rHDL. **E.** Experimental strategy used to measure macrophage phagocytosis *in vivo* based on exogenous material (DsRed) uptake. Red gate indicates the donor-derived macrophages that have incorporated host cell-derived DsRed (phagocytic macrophages). **F.** Quantitative analysis of DsRed uptake by donor-derived aortic (top left) or BM (bottom left) macrophages from DsRed chimeric mice (n = 6-7 per group) 6 hours after administration of one single dose of PBS, rHDL or PS@rHDL, measured as mean fluorescent intensity (MFI) using flow cytometry. Representative density plots of macrophages from the different groups in aorta (top right) and BM (bottom right) included in the analysis. Data was normalized to the PBS-treated group. Normally distributed data are presented as mean ± SD, and P values were calculated using one-way ANOVA, followed by multiple comparisons using Dunnett’s test. ** P < 0.01 and *** P < 0.001. MPs: macrophages.

To further explore the functional consequences of the upregulation of genes related to receptor-mediated endocytosis, we analyzed the effect of PS@rHDL on the *in vivo* phagocytic activity of macrophages in aorta and bone marrow. We used a model in which parenchymal cells but not macrophages expressed the DsRed fluorescent protein, thus allowing us to monitor macrophage phagocytic activity by flow cytometry^37,38^. We generated this model by transplanting bone marrow from SJL donor mice into ubiquitous DsRed-expressing mice^39^. This strategy allows the discrimination of donor-derived (CD45.1^+^) from recipient-derived (CD45.2^+^) macrophages to unambiguously identify transplanted macrophages that have phagocytosed extracellular-derived DsRed^+^ material (Fig. 6E). After bone marrow transplantation, macrophages (identified as CD45.1^+^ CD11b^+^ CD64^+^ MHCII^+^ cells) from SJL donors (grey, CD45.1^+^) were analyzed by flow cytometry to determine their uptake of material from DsRed^+^ cells (red). Analysis of DsRed incorporation by macrophages from aorta and bone marrow revealed increased phagocytosis in mice treated with PS@rHDL when compared to saline or rHDL (Fig. 6F). These results show that PS@rHDL enhances phagocytic activity in macrophages *in vivo*, which may underlie its beneficial effects in atherosclerotic lesions.

### PS@rHDL treatment is not immunosuppressive

Inflammation-inhibiting treatments are currently being explored as an alternative and/or complement to lipid-lowering therapies in patients at high cardiovascular risk. The CANTOS trial, testing the anti-IL-1β antibody canakinumab, afforded positive results in patients with a previous infarction. However, some fatal opportunistic infections were recorded due to the immunosuppressive action of the biological agent^8^. As an “eat-me” signal, PS has immunosuppressive effects in diverse contexts^40^, and these features have been exploited in different liposomal formulations with anti-inflammatory^41–43^ and tolerizing effects^44^. Given the observed anti-inflammatory effects of PS@rHDL, we wanted to investigate whether these were due to global immunosuppression and compare its performance against an anti-IL-1β antibody (IL-1β mAb) treatment. To that end, we carried out an infection challenge with *Staphylococcus aureus* (*S. aureus*) on WD_6w_ mice treated with saline, PS@rHDL or IL-1β mAb as shown in Fig. 7A. No differences in body mass were found among the groups after the challenge (Fig. 7B). Although the IL-1β mAb group showed the largest temperature drop and variation, the differences attained statistical significance at one time point only (Fig. 7C). However, the PS@rHDL group had a higher survival rate than the IL-1β mAb group, and no difference was found between the PS@rHDL and saline groups (Fig. 7D), indicating that PS@rHDL does not compromise the ability to fight infection when compared with unspecific inhibition of IL-1β.

**Figure 7.**
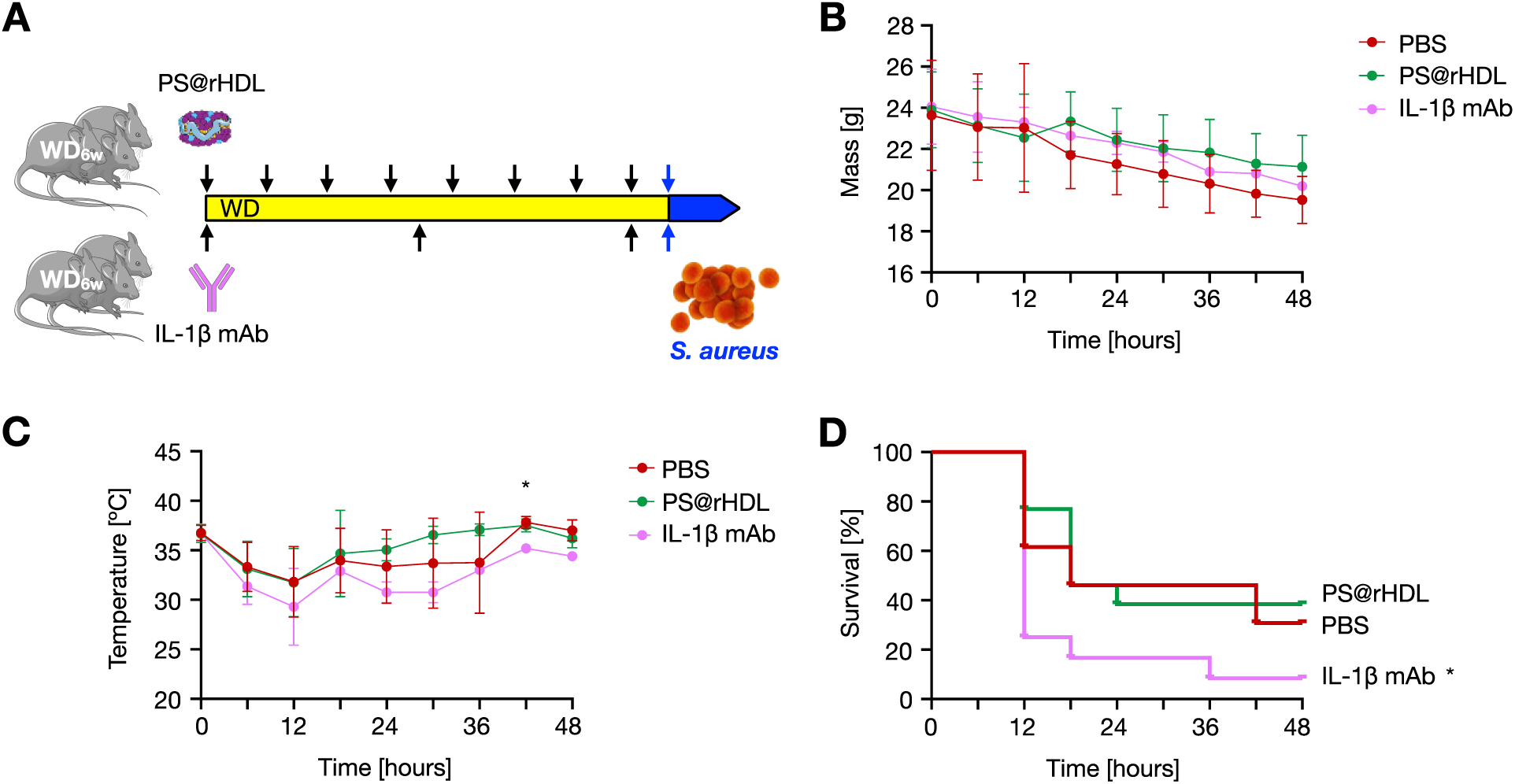
PS@rHDL treatment is not immunosuppressive. **A.** Schematic overview of the infection challenge experiment: WD_6w_ mice (n = 12-13 per group) were treated with PS@rHDL, IL-1β mAb or PBS over the course of two weeks. Subsequently mice were infected (i.p.) with *Staphylococcus aureus* (10^8^ CFU/mouse) (*S.aureus* illustration by Marina Hernández-Ávila). **B, C.** Body mass (**B**) and temperature (**C**) were monitored for 48 hours post *S. aureus* infection. **D.** Survival curves for the different treatment groups up to 48 hours post *S. aureus* infection. Data are presented as mean ± SD. P values were calculated using one-way ANOVA followed by multiple comparisons using Dunnett’s test. Gehan-Breslow-Wilcoxon test was used for comparison of survival curves. * P < 0.05. IL-1β mAb: monoclonal IL-1β antibody; WD_6w_: *Apoe^−/−^*mice fed a WD for 6 weeks. *S. aureus*: *Staphylococcus aureus*.

## DISCUSSION

Phosphatidylserines have proven anti-inflammatory effects, partly through mediating efferocytosis^22^. Our lipidomics analyses in a cohort of individuals with a heightened cardiovascular risk due to overweight or obesity revealed a strong negative association between circulating PS and the severity of carotid atherosclerotic lesions. These findings urged us to incorporate PS in reconstituted HDL nanoparticles, which efficiently target myeloid cells in the plaques^29,45^, to therapeutically exploit its anti-atherosclerosis properties in a mouse model of atherosclerosis.

In atherosclerotic lesions, PS@rHDL led to significant reductions in necrotic core size in the two disease stages investigated, namely WD_6w_ and WD_14w_. Atherosclerotic plaque necrotic cores result, in part, from impaired efferocytosis^46^, *i.e.*, clearance of apoptotic cells by macrophages and other phagocytes^22^. A reduction in necrotic core size may thus indicate improved efferocytosis by plaque macrophages, which could be due to activation of efferocytotic pathways elicited by PS, a canonical “eat-me” signal^22^. Indeed, HDL, PS and efferocytosis share a number of receptors, such as SR-BI^47^ and CD36^48,49^, and thus including PS in an HDL-like formulation such as PS@rHDL may lead to synergistic effects and augmented affinity for these receptors. *In vitro*, our formulation boosted efferocytosis compared to rHDL and free PS, and a recent study demonstrated macrophage SR-BI activation *in vitro* by PS-containing reconstituted HDL^20^. Moreover, in DsRed chimeric mice we show that PS@rHDL enhanced macrophage phagocytic activity, which added to our flow cytometry studies demonstrating PS@rHDL’s selective targeting of myeloid cells –especially macrophages in aortic tissue– strongly suggest the plausibility of this hypothesis.

A significant reduction in aortic mass was measured after PS@rHDL treatment. Given the observed moderate decrease in plaque burden, we reasoned that this mass reduction could be due to additional systemic effects on the arterial wall. In lesion-free segments of the descending aorta from both WD_6w_ and WD_14w_ mice, we observed a thickening of the arterial wall compared to WT mice. In these regions, PS@rHDL treatment normalized vessel wall area and also reduced the number of elastin fiber breaks to healthy WT control levels. Additionally, these structural changes were paralleled by a normalization in glucose uptake, which is known to be elevated at sites prior to lesion development in humans^50^, and also in dedifferentiating SMCs reliant on aerobic glycolysis^51^ as well as in activated Mo/MPs^52^. These data may be indicative of a repairing action by HDL-associated PS, preventing vessel wall remodeling before lesion formation.

Disease markers were generally improved to a greater extent in WD_6w_ compared to WD_14w_ mice after PS@rHDL treatment. A possible explanation may be the more advanced starting point in the older animals, which could potentially render their plaques less responsive to treatment. However, long-term lipid-lowering in patients also leads to a modest reduction in coronary plaque size, but results in a substantial reduction in clinical events, indicating that major changes in plaque composition and activity (necrotic core, macrophage balance, inflammation) may be more relevant to disease outcome than changes in morphology^53^. In fact, atherosclerosis regression is still poorly understood in humans^54^. In mice, however, disease regression has been reported in several studies^55,56^. Our data are compatible with certain aspects of regression, such as necrotic core size and vessel wall thickness reduction, or a shift in Mo/MP phenotype balance. Whether this rebalancing is due to a direct phenotype switch from inflammatory Mo/MPs –previously identified as a necessary condition for plaque regression in mice^57^– or to a drop in inflammatory Mo/MP numbers relative to resident-like/alternatively activated MPs through diminished recruitment and/or proliferation, remains an open question. Our results, nevertheless, do show a consistent reduction in Ly6C^hi^ monocyte and increase in CD206^+^ macrophage frequencies in all surveyed tissues, along with a substantial expansion of the resident-like/alternatively activated cluster by scRNA-seq at the expense of inflammatory and proliferating Mo/MPs. These anti-inflammatory effects of PS align well with our human *in vivo* and *in vitro* studies. In our human cohort, there was a strong inverse association between plasma PS and several inflammatory proteins with known pro-atherogenic effects, such as IL-18^58^ and resistin^59^. *In vitro*, PS@rHDL substantially reduced human primary monocyte cytokine production capacity. Moreover, inflammatory cytokine production capacity by human macrophages was lowered, also in the situation in which macrophages had been previously exposed to oxLDL^60^.

Single-cell analysis tools have enabled a detailed characterization of new cell types involved in the atherosclerotic process. scRNA-seq analysis of the aortic arch and root of WD_6w_ mice treated with either PS@rHDL or rHDL showed population changes among cell types, in addition to the rebalancing effect in the Mo/MP compartment just mentioned. We identified two clusters previously termed modulated SMCs^32^ and SEM cells/fibromyocytes^33,34^, respectively. Modulated SMCs are characterized by diminished expression of SMC canonical contractile genes and by *Lgals3* (Mac-2) expression as a transition marker^32^, whereas SEM cells express fibroblast-like ECM-remodeling (*Lum*, *Dcn*) and collagen (*Col1a1*) genes, but also *Il1r1*, *Csf1* and *Hif1a*. Both clusters, which seem to be de-differentiating transitional states toward a fibroblast phenotype, were smaller in the PS@rHDL group. This reduction could be a direct effect of the myeloid cell population rebalancing and the consequent loss of IL-1β-producing Mo/MPs and DCs. In turn, the reduced number of *Csf1*-expressing SEM cells may explain, at least in part, the drop in proliferating Mo/MPs, thus generating a feedforward loop towards resolution.

The development of mature atherosclerotic plaques takes decades, after which clinical symptoms such as MI or stroke can occur. These events themselves lead to periods of rapid acceleration of atherogenesis throughout the body, by further activation of the immune system^61,62^. Our experimental paradigm consisted in short treatments in mice with established atherosclerosis, which is a particularly relevant scenario in situations as described above. Immediately after cardiovascular events, fast-acting interventions are in great demand, and a well-defined therapeutic window for using PS@rHDL or similar nanobiologics^45^ thus opens up after MI and stroke. The inflammation ensuing these events aggravates atherosclerosis and very frequently leads to a recurring event within the following months. It is therefore paramount to stabilize the atherosclerotic plaques in these patients and prevent, at least, further disease progression. This was the rationale behind the CANTOS trial^8^, testing canakinumab (anti-IL-1β mAb) in patients with a history of MI. The outcomes were positive but canakinumab treatment resulted in an unacceptably high number of deaths due to opportunistic infections. In this study, we explored this possibility for PS@rHDL, since PS could potentially cause similar detrimental immunosuppressive effects^40^. Compared to an anti-IL-1β mAb, PS@rHDL showed reduced mortality upon an infectious challenge with *S. aureus*, while there was no difference with the saline control group. This suggests that HDL-associated PS does not elicit detrimental suppression of immune responses^40^.

Some limitations of our study need to be mentioned. Firstly, we could not perform lipoprotein-based lipidomics analysis of the human samples and thus we cannot ascertain in which fraction PS is more enriched. However, the significant correlations with HDL-cholesterol, but not TGs or LDLs, strongly suggest enrichment in HDL. Secondly, the PS-free rHDL formulation we used as control cannot be deemed as proper HDL, as it only contains the apolipoprotein and DMPC. HDL has a more complex composition in its many forms and their atheroprotective activity has not been directly compared to our control formulation. Finally, here we used the *Apoe^-/-^* mouse model of atherosclerosis, which is just one of the available models. In a recent study using a PS-enriched HDL formulation in *Ldlr^-/-^* mice, another widely used model of the disease, the authors did not observe a reduction in plaque size^20^. This could well be due to the different model used, but also to differences in formulation composition, dosing or treatment regimen.

In conclusion, our results are compatible with HDL-associated PS having both inflammation-resolving and plaque-stabilizing effects. The failure of HDL-raising therapies has shifted the research focus toward improving/enhancing HDL’s function rather than just augmenting circulating concentrations^15,63^. Due to its inherent biocompatibility, formulation of PS into HDL-like compositions thus holds promise as a feasible approach to potentiate their atheroprotective functions with a view to clinical translation.

## Acknowledgements

The authors thank CNIC core facilities (flow cytometry, histopathology) as well as the advance imaging and comparative medicine units for their assistance. We also thank Dr. Beatriz Salinas for kindly providing *Staphylococcus aureus*, and Dr. David Sancho for extensive discussion of results and feedback on the manuscript. Parts of the figures were drawn by using pictures from Servier Medical Art by Servier, licensed under a Creative Commons Attribution 3.0 Unported License (https://creativecommons.org/licenses/by/3.0/), and BioRender.

## Author contributions

Specific author contributions are as follows: C.P-M. and N.P.R. conceptualized the study. A.B., A.H., C.G-R., A.S., J.A.N.-A., C.T., A.D., F.S-C., K.P., C.Th., M.G.N., W.J.M.M., N.P.R. and C.P-M. engaged in the conception and design of experiments. A.B., A.H., C.G-R, I.S., A.S., J.A.N.-A., L.P.-C., M.Y., Al.B., A.R., M.L.S., and M.M.T.v.L performed the experiments and participated in the data acquisition process. A.B., A.H., C.G-R., A.S., J.A.N.-A., L.P.-C., M.Y., C.T., L.S-G., F.S-C., A.H., M.L.S, M.M.T.v.L, L.S, J.H., K.P., C.Th., M.G.N., W.J.M.M., N.P.R. and C.P-M. performed the analysis and interpretation of the data. A.B., A.H., N.P.R. and C.P-M. drafted the manuscript. All authors edited the manuscript and agreed on its final version.

