## Supplementary material for "HDL-associated phosphatidylserine blunts myeloid activation and protects from atherosclerosis": Supplemmentary material

### -Supplementary Material File-

#### METHOD DETAILS

##### Human lipidomics

***Cardiovascular measurements.*** Cardiovascular phenotyping of the cohort encompassed detailed carotid intima-media thickness (IMT) evaluations, carotid plaque detection, and assessments of plaque thickness across various carotid artery segments. Preparation involved overnight fasting or a 6-hour post-breakfast wait, along with a 12-hour abstinence from caffeine and smoking. Measurements were conducted in a temperature-regulated environment, with participants in a supine position. A 7.5-MHz transducer on a Mylab class C ultrasound device was employed to assess the baseline diameter, distensibility, and wall thickness of the carotid artery, focusing on the proximal 1-cm straight portion from three angles during diastole. Carotid IMT, an essential measure of cardiovascular health reflecting the combined thickness of the inner two layers of the carotid artery wall, was determined using an automated boundary detection system.

Additionally, the study measured the presence and maximum thickness of plaques in the common, internal, and external carotid arteries, and at the carotid bulbous. Plaque presence was identified by focal wall thickening at least 1.5 times the mean IMT or an IMT exceeding 1.5 mm, aligning with the Mannheim IMT consensus[1]. Body fat distribution was evaluated through magnetic resonance imaging (MRI), categorizing volumes of visceral adipose tissue and subcutaneous adipose tissue into deep and superficial compartments. Hepatic fat content was quantified utilizing localized proton magnetic resonance spectroscopy, providing comprehensive insights into the participants' cardiovascular and metabolic health profiles[2,3].

***Metabolic parameters.*** The metabolic syndrome (MetS) was defined according to the National Cholesterol Education Program Adult Treatment Panel III criteria[4] as a minimum of three listed features: 1) Abdominal obesity is characterized by a waist circumference exceeding 102 centimeters (40 inches) in males and 88 centimeters (35 inches) in females. 2) Triglyceride levels equal to or greater than 150 mg/dL (1.7 mmol/L), or receiving pharmacological intervention for elevated triglycerides. 3) High-density lipoprotein (HDL) cholesterol levels below 40 mg/dL (1.0 mmol/L) in males and below 50 mg/dL (1.3 mmol/L) in females, or undergoing treatment for reduced HDL cholesterol. 4) Blood pressure readings equal to or exceeding 130/85 mmHg, or receiving pharmacological treatment for hypertension. 5) Fasting plasma glucose levels equal to or greater than 100 mg/dL (5.6 mmol/L), or undergoing treatment for hyperglycemia. Serum insulin and glucose levels were measured and used to calculate insulin resistance employing the homeostatic model assessment for insulin resistance (HOMA IR)[5].

***Measurement of circulating lipids.*** Biochemical parameters were assessed following established protocols[6]. Morning blood samples after overnight fasting measured glucose, total cholesterol (TC), triglycerides (TG), and HDL cholesterol (HDL-C) through standard lab methods, while LDL cholesterol (LDL-C) was calculated via the Friedewald formula. Fresh whole blood EDTA samples were analyzed for immune cell counts using the Sysmex XE-5000 (Syxmex, XE-5000). Circulating cytokines and proteins, including high-sensitivity C-reactive protein (R&D Systems, DY1707), interleukin (IL)-6 (Sanquin, M9316), IL-18-bp (Protein Simple, SPCKB-PS-000501), vascular endothelial growth factor (VEGF) (Protein

Simple, SPCKB-PS-000330), alpha-1 antitrypsin (AAT) (R&D Systems, DY1268), resistin (R&D Systems, DY1359), leptin (R&D Systems, DY398), adiponectin (R&D Systems, DY1065), and IL-18 (Protein simple, SPCKB-PS-000501), were quantified using ELISA kits.

***Peripheral Blood Cell Isolation and Stimulation.*** Blood from participants was collected into EDTA tubes (Monoject) to isolate PBMCs using a modified protocol: dilution in phosphate-buffered saline (PBS), density centrifugation over Ficoll-Paque (GE Healthcare), then washing and resuspension in RPMI 1640 medium (Invitrogen, 11875093) supplemented with 50 µg/mL gentamicin (Centrafarm), 2 mM GlutaMAX™ (Life Technologies, 35050061), and 1 mM sodium pyruvate (Life Technologies, 11360070). Red blood cells were lysed using Zap-Oglobin II (Beckman Coulter, 7546138), and mononuclear cells, adjusted to 5 × $10^5$ cells/100 µL, were placed into 96-well plates[7], and stimulated with 1 ng/mL lipopolysaccharide (LPS) (Sigma-Aldrich; from *Escherichia coli* serotype 055:B5, Sigma-Aldrich, L2880) and 1 µg/mL Pam3Cys-Ser-(Lys)4 (P3C) (EMC Microcollections; L2000) for 24 hours. Supernatants were then collected for cytokine production assessment of IL-1β (R&D Systems, DY201), TNF-α (R&D Systems, DY210), IL-1Ra (R&D Systems, DRA00B), and IL-6 (Sanquin, M9316) by ELISA[6].

**Lipidomic Analysis**

***Chemical Solvent and Reagent Quality Assurance.*** All solvents used were of high-performance liquid chromatography-mass spectrometry grade (HPLC-MS) grade (VWR International GmbH and Merck KGaA), to ensure the reliability and accuracy of our lipidomic analyses, as underscored in previous studies[8]. The internal standard mixtures was manually prepared for 2ul of plasma samples and consisted of 210 pmol PE(31:1), 396 pmol PC(31:1), 99 pmol PS(31:1), 85 pmol PI(34:0), 56 pmol PA(31:1), 52 pmol PG(28:0), 29 pmol CL(56:0), 40 pmol LPA(17:0), 35 pmol LPC(17:1), 38 pmol LPE(17:0), 32 pmol Cer(17:0), 99 pmol SM(17:0), 55 pmol GlcCer(12:0), 37 pmol GM3(18:0-D3), 340 pmol TG(50:1-d4), 740 pmol CE(17:1), 64 pmol DG(31:1), 104 pmol MG(17:1), 724 pmol Chol(d6), 45.99 pmol Car(15:0).

***Sample preparation.*** The content of lipids in the mass spectrum was quantified in the mass spectrometer-Thermo Q Exactive Plus. The lipids were extracted using 500 µL of 1/5 of chloroform/methanol that contained the internal standard mix. The samples were then sonicated for a minute with an ultra-sonic bath and centrifuged at 20,000 × g for 5 minutes. The supernatant was transferred into a fresh tube, 200 µL chlorofom and 600 µL of 1% acetic acid were added followed by brief vortexing and centrifugation at 20,000 × g for 2 minutes[9]. The aqueous layer was aspirated and the organic phase was transferred to a fresh tube and then evaporated in a Speed Vacuum Concentrator set at 45°C for 20 min. In the end, the lipid was reconstituted in 350 µL of a spray buffer (isopropanol/methanol/water 8/5/1, 10 mM ammonium acetate) and sonicated for 5 minutes to have a homogeneous solution[10]. For quality control, each batch contained positive and negative controls for any potential evidence of batch effects.

***Instrumental Configuration.*** Samples were analyzed on the Thermo Q Exactive Plus mass spectrometer, equipped with a heated electrospray ionization II (HESI II) source. This setup was specifically chosen for its proven efficacy in facilitating shotgun lipidomics, allowing for the detailed and precise profiling of lipid species[11,12].

***Data Acquisition.*** The mass spectrometer was calibrated to perform high resolution at 280,000 to collect MS1 spectra over a mass-to-charge (m/z) range of 250 to 1200 using the positive ionization mode. MS/MS spectra were acquired using the data-independent acquisition at a resolution of 70,000 in 1 m/z windows from 250 to 1200 m/z using the positive ionization mode. The final coverage of the lipidomic analysis arises from the entire sum of the two modes of acquisition.

***Data Processing and Analysis.*** Mass spectrometry .raw were converted into .mzML and analyzed using the LipidXplorer software[13]. Lipid species were identified with the aid of custom MFQL scripts and quantified against internal standards intensity. The lipidomic profiles obtained were normalized against the total extracted plasma volume, thus allowing for solid comparative analyses to be done across diverse samples[14].

**Human *in vitro* procedures**

***Human PBMC and monocyte isolation.*** Human PBMCs were isolated by Pancoll (PAN-Biotech, P04-601000) and monocytes by a hyper-osmotic Percoll (Sigma-Aldrich, P1644) density gradient centrifugation. The cells were resuspended and cultured in RPMI 1640 Dutch modified medium (Thermo Scientific, 22409031), supplemented with 5 µg/mL gentamicin (Thermo Scientific, 15710064), 2 mM Glutamax (Thermo Scientific, 35050061), 1 mM sodium pyruvate (Thermo Scientific, 11360070). PBMCs were pre-treated with RPMI or with different concentrations of free-PS, rHDL, and PS@rHDL (50 µg/mL and 200 µg/mL) and stimulated with 10 µg/mL Pam3Cys-Ser-(Lys)4 (Pam3Cys) (EMC Microcollections; L2000) or 10 ng/mL LPS (Sigma-Aldrich; from *Escherichia coli* serotype 055:B5, Sigma-Aldrich, L2880) for 24h.

***PBMCs and monocytes* in vitro *procedures.*** The *in vitro* protocol was followed as previously described[15]. Briefly, Percoll-isolated monocytes were left to adhere to polystyrene flat-bottom plates (Corning, CLS3599) for 1 h at 37°C 5% $CO_2$ and washed with warm PBS, monocytes were incubated with media only or with 10 µg/mL oxLDL (prepared from human isolated LDL as previously described[16] for 24 h in the presence of 10% human pooled serum). The culture medium was refreshed on day 3. At day 6 the cells were incubated with the media as control or pretreated for 30 minutes with free-PS, rHDL or PS@rHDL at concentrations of 50 ug/mL or 200 ug/mL and then re-stimulated for another 24h with 10 µg/mL Pam3Cys-Ser-(Lys)4 (Pam3Cys) (EMC Microcollections; L2000) or 10 ng/mL LPS (Sigma-Aldrich; from *Escherichia coli* serotype 055:B5, Sigma-Aldrich, L2880) for 24h.

***Cytokine quantification.*** IL-6 and TNF-α production in supernatants was determined using commercial IL-6 and TNF-α DuoSet ELISA kits (R&D Systems, DY210), according to the instructions of the manufacturer.

**Nanoformulation**

***Chemicals.*** Apolipoprotein A1 (apoA1) was isolated from HDL from human plasma (Bioresource Technology, H3025) and purified as previously described[17]. The phospholipids

1,2-dimyristoyl-sn-glycero-3-phosphocholine (DMPC, 850345), 1,2-distearoyl-sn-glycero-3-phosphoethanolamine (DSPE, 850715), and phosphatidylserine (PS, 840032) were purchased from Avanti Polar Lipids. 1-(4-Isothiocyanatophenyl)-3-[6,17-dihyroxy-7,10,18,21-tetraoxo- 27-[N-acetylhydroxyla-mino) 6,11,17,22-tetraazaheptaeicosane] thiourea (DFOp-NCS) was purchased from Macrocyclics. The phospholipid chelator DSPE-DFO was synthesized from DSPE and DFOp-NCS as reported. The fluorescent label 1,10-Dioctadecyl-3,3,30,30-Tetramethylindotricarbocyanine Iodide (DiR) was purchased from AAT Bioquest (22070). All other reagents were acquired from Sigma-Aldrich/Merck unless otherwise stated.

***Synthesis and characterization of apoA1 formulations.*** ApoA1 formulations were prepared following previously published reconstitution methods[18]. Briefly, for the PS-containing apoA1 formulation, henceforth PS@rHDL, DMPC and PS were mixed in a 4:1 mass ratio in chloroform and then dried in vacuum. The resulting lipid film was hydrated with a phosphate buffered saline (PBS) solution of apoA1 at a 1:5 apoA1/phospholipid mass ratio. The mixture was then sonicated for 30 minutes using a Sonoplus HD 2070 tip ultrasonic homogenizer (Bandelin) working at 65-70% power output. Subsequently, the solution was centrifuged at 5,000 rpm for 10 minutes and filtered through a 0.22 µm PES filter. The PS-free apoA1 formulation, namely rHDL, was prepared following the same procedure but using only DMPC (5:1 DMPC/apoA1 mass ratio). ApoA1 concentration was quantified using a BCA test (ThermoFisher Scientific, 23225), and PS concentration was determined using a fluorimetric assay kit (Sigma-Aldrich, MAK371). For targeting, pharmacokinetics and biodistribution experiments, labeled analogues of PS@rHDL were prepared through incorporation in the initial phospholipid mix of the fluorescent dye DiR (1% of total lipids), or the phospholipid chelator DSPE-DFO[19] (1% at the expense of DMPC), which enables radiolabeling with $^{89}$Zr[18]. All formulations were characterized by a combination of dynamic light scattering, zeta potential and size exclusion chromatography (SEC) measurements. SEC was performed on a Jasco 1200 high-performance liquid chromatography (HPLC) system equipped with two LC-10AT pumps and an SPD-M10AVP photodiode array detector, and a Superose 6 increase 10/300 GL column (Cytiva, 29-0915-96) using PBS as eluent at a flow rate of 0.5 or 0.75 mL/min. Hydrodynamic size and z-potential were measured on a Zetasizer Nano ZS90 (Malvern).

***Radiolabeling of PS@rHDL with $^{89}$Zr.*** $^{89}$Zr was supplied by Perkin Elmer, and activity measurements were made using a AtomLab500 dose calibrator (Biodex). A solution of 1% DSPE-DFO-bearing PS@rHDL in PBS was reacted with $^{89}$Zr-oxalate at 37ºC for 1 hour[18]. The labeled nanoparticles were separated from free unreacted $^{89}$Zr by gel filtration using a PD-10 column (Cytiva, 17085101) and PBS as eluent. The radiochemical yield was 73±2% (n=2) and the radiochemical purity >97%.

#### *In vitro* murine procedures

***Cholesterol efflux assay.*** Bone marrow cells were flushed from femurs of 4-month-old C57BL/6J mice and differentiated into BMDMs by incubation in RPMI 1640 media (Gibco, 21875-034) supplemented with 10% fetal bovine serum (FBS) (Sigma-Aldrich, F7524-1654682), 1% penicillin (10000U/mL) / streptomycin (10mg/mL) (P/S) (Sigma-Aldrich, P4333), and 30% L929 cell-conditioned media (as a source of M-CSF) for 7 days. BMDMs were washed twice with PBS and equilibrated for 18 hours in media containing RPMI, 1% P/S, 30% L929-conditioned medium and 2 mg/mL bovine serum albumin (BSA) (VWR, A7906-10G). Cells were subsequently incubated with media containing 0.5 µCi/mL [$^3$H]-cholesterol (PerkinElmer, NET139250UC) and 50 µg/mL acetylated LDL (Invitrogen, L35354) for 24 hours. Cells were washed twice with PBS and incubated with cholesterol acceptors (rHDL (50 µg/mL), PS@rHDL (50 µg/mL) and PS (50 µg/mL)) and PBS as control for 8 hours. Culture media was harvested and cleared of cellular debris by brief centrifugation, and cells were lysed in 0.1 M NaOH solution for 2 hours. The [$^3$H]-cholesterol content of media and cell lysates was measured in a LS 6500 Multi-Purpose Scintillation Counter (Beckman Counter). Efflux is expressed as % of efflux over control:

$$\%\ efflux = \left(\frac{cpm\ in\ medium}{cpm\ in\ medium + cpm\ in\ cells} * 100\right)_{rHDL\ or\ PS@rHDL\ or\ PS} - \left(\frac{cpm\ in\ medium}{cpm\ in\ medium + cpm\ in\ cells} * 100\right)_{PBS}$$

***Efferocytosis* in vitro.** Bone marrow cells were flushed from femurs of 4-month-old C57BL/6J mice and differentiated into BMDMs by incubation in RPMI 1640 media (Gibco, 21875-034) supplemented with 10% fetal bovine serum (FBS) (Sigma-Aldrich, F7524-1654682), 1% penicillin (10000U/mL) / streptomycin (10mg/mL) (P/S) (Sigma-Aldrich, P4333), and 30% L929 cell-conditioned media (as a source of M-CSF) for 7 days. Thymocytes isolated from male C57BL/6J 5 weeks aged mouse were labeled with Cell Trace Violet stain (ThermoFisher Scientific, C34557) in PBS for 20 minutes. The cells were exposed to ultraviolet for apoptosis induction for 10 minutes and cultured for 2 hours. Then Trace Cell Violet positive and apoptotic thymocytes were cocultured with BMDM (5:1) pretreated with PBS, rHDL (50 µg/mL), PS@rHDL (50 µg/mL), or PS (50 µg/mL) for 24h. Two hours later thymocytes were washed out and BMDM were stained with anti-CD11b (clone M1/70)-APC (RRID: AB_312795, Cat#101212, BioLegend) and anti-F4/80 (clone BM8)-A488 (RRID: AB_469915, Cat#53-4801-82, eBioscience) for macrophages selection, and zombie yellow (BioLegend, 423104) for viability selection. Cell violet fluorescence was used for apoptotic engulfed thymocytes quantification. Samples were analyzed using a BD LSRFortessaTM cytometer (BD Biosciences) equipped with BD FACSDiva™ software version 6.2 (BD Biosciences) and the resulting data were analyzed using FlowJo software version 10.9.

#### Animal procedures

***Pharmacokinetics.*** $WD_{6w}$ and $WD_{14w}$ mice (n = 4-5 per group) were injected with $^{89}$Zr@PS@rHDL (10-15 µCi, ~20 mg apoA1/PS/kg). At predetermined time points (2, 10, 30 minutes and 1, 4, 8, 24 hours) blood was collected, weighed and gamma counted using a Wizard 1470 automatic gamma counter (PerkinElmer). Data were corrected for decay, converted to percentage of injected dose per gram of tissue (%ID/g), plotted in a time-activity curve and fitted using a non-linear two-phase decay regression in GraphPad Prism software, version 9.4.1. A weighted blood radioactivity half-life ($t_{1/2w}$) was finally calculated as follows:

$$t_{1/2w} = X_{fast} * t_{1/2fast} + X_{slow} * t_{1/2slow}$$

***Biodistribution.*** $WD_{6w}$ and $WD_{14w}$ mice (n = 4-5 per group) were injected with $^{89}$Zr@PS@rHDL (10-15 µCi, ~20 mg apoA1/PS/kg). Radioactivity biodistribution was measured at 24 hours post $^{89}$Zr@PS@rHDL injection. Following euthanasia, mice were perfused with ample PBS and tissues of interest collected (bone marrow, liver, spleen, kidney, aorta, muscle, heart, lungs, blood, plasma, blood cells and brain), weighed and gamma counted. Results were corrected for decay, and radioactivity concentration was normalized to tissue weight, reporting data as %ID/g.

***Targeting assay.*** $WD_{6w}$ and $WD_{14w}$ mice (n = 4 per group) were injected intravenously via the lateral tail vein with the corresponding PS@rHDL fluorescent analogue incorporating DiR, a lipophilic fluorophore. DiR-labeled PS@rHDL (~0,5 mg DiR/kg) was allowed to circulate for 24 hours. Aorta, blood, bone marrow and spleen were collected, processed to obtain a single-cell suspension and analyzed by flow cytometry (see *Flow Cytometry* section for details). The same flow cytometry settings were applied to all tissues. CD206-positive macrophages were not discriminated. DiR was detected on the APC-Cy7 channel. A negative control, *i.e.*, a mouse not injected with DiR-labeled PS@rHDL, was used to define $DiR^+$ gate. All data were acquired using a BD LSRFortessaTM cytometer (BD Biosciences) equipped with BD FACSDiva™ Software v 6.2 (BD Biosciences) and analyzed using FlowJo v10.9 software.

***Treatment schedule.*** $WD_{6w}$ and $WD_{14w}$ mice were randomly allocated to four treatment groups, receiving PBS (intravenous (i.v.), 100 µL); rHDL (~20 mg apoA1/kg, i.v.); PS@rHDL (20 mg PS/kg, i.v.); or PS (20 mg/kg, intraperitoneal). Intravenous doses were administered via lateral tail vein injection. The treatment schedule consisted of 8 administrations over two weeks (see Fig. 3A), during which time mice were kept on WD. Animals were fasted from the time of last administration until euthanasia 24 hours later, following which they were perfused with PBS and tissues of interest collected and weighed.

***Toxicity.*** Potential adverse effects were assessed through a complete blood count hematological analysis, performed using an ABX Pentra 80 hematology analyzer (Horiba Medical v1.12.0). Additionally, a lipid profile was obtained from plasma samples using a Dimension RxL Max System (Siemens).

***$^{18}$F-FDG study.*** After the last treatment injection, mice were fasted for 24 hours before [$^{18}$F]-2-deoxy-2-fluoro-D-glucose ($^{18}$F-FDG, ~300 µCi, Curium Pharma) administration. $^{18}$F-FDG was administered via a lateral tail vein and allowed to circulate for 1 hour. Mice were anesthetized with isofluorane (Ecuphar, IsoFlo, 100% p/p), euthanatized and perfused with PBS for *ex vivo* analysis. Aorta, blood, liver, spleen, bone marrow and muscle were collected, blotted, weighed and counted on a Wizard 1470 automatic gamma counter. Results were corrected for decay, and radioactivity concentration was normalized to tissue weight, reporting data as percentage of injected dose (%ID) or as percentage of injected dose per gram of tissue (%ID/g).

**In vivo *phagocytic activity quantification.*** For generation of transplanted mice, recipient DsRed mice[20] were lethally irradiated (6.5 Gy split doses, 3 hours apart), and subsequently received BL/6.SJLC57 (CD45.1) BM cells. Donor BM cells were harvested by flushing both femurs into RPMI and one million were intravenously injected into irradiated recipients. Animals were analyzed 6-8 weeks after BM transplantation. Recipient mice were injected intravenously once with PBS, rHDL (20 mg/kg), or PS@rHDL (20 mg/kg) and 6 hours later tissues were harvested. For identification of phagocytic macrophages, single-cell suspensions were obtained from BM through mechanical dissociation by flushing and from aorta as previously described in the "Flow cytometry" section. Phagocytic macrophages (MPs) were identified based on the incorporation of DsRed fluorescence using the following antibodies: anti-CD45.1 (clone A20)-PerCP.Cy5.5 (Cat#65-0453, TonboBioscience), anti-CD11b (clone M1/70)-Bv510 (RRID: AB_2561390, Cat#101245, Biolegend), anti-CD64 (clone X54-5/7.1)-APC (RRID: AB_11219391, Cat#139306, Biolegend) and anti-MHCII (clone M5/114.15.2)- PE.Cy7 (RRID: AB_2069376, Cat#107630, Biolegend). Engulfment of material in bone marrow chimeras was based on acquisition of DsRed signal by CD45.1-derived macrophages as compared to non-fluorescent (WT) control mice.

***Infection challenge experiment.*** $WD_{6w}$ mice were randomly assigned in three treatment groups (n = 12-13 per group) and injected intravenously via lateral tail vein with PBS, PS@rHDL (standard treatment at 20 mg/kg), or a monoclonal IL-1β antibody (3 injections at 10 mg/kg, RRID: AB_2687727, Abyntek BioPharma, BE0246-5MG). Twenty-four hours after the last injection, mice were infected with *Staphylococcus aureus* (~$10^8$ CFU/mouse) intraperitoneally. Body mass, temperature and survival were monitored over the following 48 hours.

**_Ex vivo_ procedures**

***Flow cytometry.*** Aorta, bone marrow and spleen samples were processed to obtain a single-cell suspension. After adipose tissue removal, aortas were diced and digested with a cocktail of enzymes, including liberase TH (4 U/mL; Sigma-Aldrich, 5401151001), hyaluronidase type I-S (60 U/mL; Sigma-Aldrich, H3506) and deoxyribonuclease I (40 U/mL; Sigma-Aldrich, DN25) in PBS at 37ºC for 1 hour in an Eppendorf ThermoMixer C thermoblock shaking at 300 rpm. Cells were resuspended in 2% fetal bovine serum (FBS) PBS and passed through 70-µm cell strainers (VWR, 734-2761) to obtain single-cell suspensions prior to antibody staining. Surrounding tissue from femurs and tibias was removed and bones were crushed in cold PBS using a mortar and pestle. The solution was filtered through a 40-µm cell strainer (VWR, 732-2760). Spleens were smashed and the solution was filtered with a 40-µm cell strainer. Red blood cells from single-cell suspensions of bone marrow and spleen, and from blood (100 µL) were lysed with 0.15 M ammonium chloride (Sigma-Aldrich, A9434-500G) for 10 minutes. Single cell suspensions from aorta, bone marrow, blood and spleen thus prepared were immunophenotyped using the antibodies the following antibodies: anti-CD90.2 (clone 53-2.1)-Biotin (RRID: AB_10643274, Cat#140314, BioLegend), anti-CD49b (clone DX5)-Biotin (RRID: AB_313411, Cat#108904, BioLegend), anti-B220 (clone RA3-6B2)-Biotin (RRID: AB_312989, Cat#103204, BioLegend), anti-Ter119 (clone TER-119)-Biotin (RRID: AB_313705, Cat#116204, BioLegend), anti-Ly6G (clone 1A8)-Biotin (RRID: AB_1186108, Cat#127604, BioLegend,) and NK1.1 (clone PK136)-Biotin (RRID: AB_313391, Cat#108704, BioLegend) to identify T cells, B cells, erythrocytes, granulocytes and natural killer cells respectively, identified in bulk as a dump gate for non-myeloid immune cells. To characterize myeloid cells, dendritic cells, monocytes, and macrophages, antibodies recognizing anti-CD11b (clone M1/70)-APC (RRID: AB_312795, Cat#101212, BioLegend), anti-CD11c (clone N418)-PerCP Cy5.5 (RRID: AB_2129641, Cat#117328, BioLegend), anti-Ly6C (clone AL-21)-FITC (RRID: AB_394628, Cat#553104, BD Pharmingen), anti-F4/80 (clone BM8)-PE.Cy7 (RRID: AB_893478, Cat#123114, BioLegend) and anti-CD206/MMR (clone MR6F3)-

APC-eFluor780 (RRID:AB_2802285, Cat#47-2061-82, eBioscience) were used. Subsequently single-cell suspensions were incubated with streptavidin-phycoerythrin (BioLegend, 405204). Cell viability was assessed using 4',6-diamidino-2-phenylindole dihydrochloride (DAPI) (Sigma-Aldrich, D8417). Samples were analyzed using a BD LSRFortessaTM cytometer (BD Biosciences) equipped with BD FACSDiva™ software version 6.2 (BD Biosciences) and the resulting data were analyzed using FlowJo software version 10.9.

***Histology and immunohistochemistry.*** Aortas were fixed in 4% paraformaldehyde (VWR, 43368.9M) for 48 hours and subsequently surrounding adipose tissue was removed. The thoracic aorta was divided in root, ascending aorta, brachiocephalic artery bifurcation and descending aorta. Samples were paraffined and sectioned into 4 µm serial sections. Root, ascending aorta and bifurcation sections were stained with Masson's Trichrome to quantify plaque, necrotic core, tunica media, and collagen areas. Root sections were also stained with Alizarin red to quantify calcifications. Immunohistochemistry was performed using Mac-2 antibody (1:800) (polyclonal rat anti-Mac-2 (clone M3/38), RRID: AB_837132, Cat#14-5301-8, Invitrogen) and secondary antibody rabbit anti-rat HRP (RRID: AB_228439; Cat#ab6734, Abcam) to quantify inflammatory macrophage and foam cell burden in root sections. Descending aortas were analyzed at two different levels, namely at the end of the aortic arch meeting the beginning of the descending aorta (upper level, see Fig. 4C), and at the level of the diaphragm (lower level, see Fig. 4C). Slides were stained with hematoxylin and eosin (H&E) for media thickness and nuclei quantification; with Masson's Trichrome (MT) for collagen content quantification; and with Van Gieson's stain (VG) to quantify elastin fiber breaks. Slides were digitalized using an AxioSacan Z1 (Carl Zeiss). Images were visualized using NDPview 2 software, and quantifications were performed using ImageJ software. Normalized parameters were calculated versus total intima and media area.

***Single-cell RNA sequencing (scRNA-seq).*** Aortic arch and root samples from $WD_{6w}$ mice were collected after treatment with either rHDL or PS@rHDL (n = 6 per group). Mice were fasted for 24 hours and following euthanasia were perfused with 5mM EDTA PBS (Sigma-Aldrich, E1644). After adipose tissue removal, aortic arch and root samples were diced and digested with a cocktail of enzymes to prepare a single-cell suspension as described above (see Flow cytometry). Single-cell suspensions were pooled into one sample per treatment group. Cells were then incubated with DAPI (Sigma-Aldrich, D8417) and DRAQ5 (Invitrogen, 65-0880-96) for dead cell exclusion. Cells were sorted using a FACS Aria II cell sorter equipped with BD FACSDiva™ Software v6.1.3. Cells were counted and their viability was checked using the Countess 3 cell counter (ThermoFisher). Each cell suspension was loaded into one port of a Chromium Next GEM Chip G (10x Genomics) with a target output of 10,000 cells. Single cells were encapsulated into emulsion droplets using the Chromium Controller (10x Genomics). scRNA-seq libraries were prepared using the Chromium Next GEM Single Cell 3' Kit v3.1 (10x Genomics) following the manufacturer instructions and each library was amplified using a SureCycler 8800 thermal cycler (Agilent Technologies). The average size of each library was then calculated using a High sensitivity DNA chip on a 2100 Bioanalyzer (Agilent Technologies) and the concentration was determined using the Qubit fluorometer (ThermoFisher). Individual libraries were diluted to 10 nM and pooled for sequencing. Library pool was loaded at 650 pM onto a P3 flow cell (100 cycles) of the NextSeq 2000 (Illumina) in paired-end configuration (28bp Read1, 10bp Index1, 10pb Index 2 and 90bp Read2). FastQ files for each sample were obtained using cellranger mkfastq pipeline (10x Genomics).

***scRNA-seq bioinformatics analysis.*** Single cell identification and transcriptome profile has been obtained using the 10X CellRanger pipeline (v6.1.1)[21] and mouse mm10 genome reference (GENCODE vM23/Ensembl 98). CellBender software (v0.2.2)[22] was used to eliminate technical artifacts from the high-throughput single-cell RNA sequencing data with the following parameters: –expected-cells 36020, --total-droplets-included 38020, --fpr 0.01 and 150 epochs. Single Cell filtering and clustering have been done using Scater[23] and Seurat[24] R packages. Cells have been filtered by a sequencing depth between 800 and 45000, a minimum of 2000 genes detected, a mitochondrial content below 15%, a cell fraction above 0.2, a gene expression complexity below 60% (to filter out cells with high levels of reads in just a few genes) and a hemoglobin gene set expression below 0.1%. At the end of the filtering process, a total of 12721 cells were retained, that were log-normalised and scaled. Doublets were identified using scDblFinder (Version v1.12.0)[25]. Clustering and dimensionality reduction of these cells were performed using Seurat R methodology with the 2000 most variable genes and 18 PCs. Cells in clusters with high density of doublets or high levels of background RNA have been removed and reclustered again using the 2000 most variable genes and 18 PCs. Cluster’s markers and differential expression analysis between conditions have been perform using MAST algorithm and testing only genes detected in 30% for markers and 10% for differential expression on any cluster. A functional analysis was performed using enrichR[26,27], to identify enriched pathways between conditions. Myeloid clusters were subset (1010 cells) and reclustered using the 2000 most variable genes and 20 PCs. Cluster’s markers, differential expression analysis between conditions and enrichment analysis were performed as before.

| REAGENTS | SOURCE | IDENTIFIER |
|---|---|---|
| **Antibodies** | | |
| Anti-CD90.2-Biotin (clone 53-2.1) | BioLegend | Cat#140314; RRID: AB_10643274 |
| Anti-CD49b-Biotin (clone DX5) | BioLegend | Cat#108904; RRID: AB_313411 |
| Anti-B220-Biotin (clone RA3-6B2) | BioLegend | Cat#103204; RRID: AB_312989 |
| Anti-Ter119-Biotin (clone TER119) | BioLegend | Cat#116204; RRID: AB_313705 |
| Anti-NK1.1-Biotin (clone PK136) | BioLegend | Cat#108704; RRID: AB_313391 |
| Anti-Ly6G-Biotin (clone 1A8) | BioLegend | Cat#127604; RRID: AB_1186108 |
| Anti-CD45.1-PerCP.Cy5.5 (clone A20) | TonboBioscience | Cat#65-0453 |
| Anti-CD11b-APC (clone m1/70) | BioLegend | Cat#101212; RRID: AB_312795 |
| Anti-CD11b-Bv510 (clone M1/70) | Biolegend | Cat#101245; RRID: AB_2561390 |
| Anti-CD64-APC (clone X54-5/7.1) | Biolegend | Cat#139306; RRID: AB_11219391 |
| Anti-MHCII-PE.Cy7 (clone M5/114.15.2) | Biolegend | Cat#107630; RRID: AB_2069376 |
| Anti-CD11c- PerCPCy5.5 (clone N418) | BioLegend | Cat#117328; RRID: AB_2129641 |
| Anti-Ly6C-FITC (clone AL-21) | BD Pharmingen | Cat#553104; RRID: AB_394628 |
| Anti-F4/80-PE.Cy7 (clone BM8) | BioLegend | Cat#123114; RRID: AB_893478 |
| Anti-CD206- APCeF780 (clone MR6F3) | eBioscience | Cat#47-2061-82; RRID: AB_2802285 |
| Anti-Mac-2 (clone M3/38) | Invitrogen | Cat#14-5301-85; RRID: AB_837132 |
| rabbit anti-rat-HRP (IgG H&L) | Abcam | Cat#ab6734; RRID: AB_228439 |
| Monoclonal anti-IL-1β (clone B122) | Abyntek BioPharma | Cat#BE0246, RRID: AB_2687727 |
| **Chemicals** | | |
| 1,2-dimyristoyl-sn-glycero-3-phosphocholine, powder, 14:0 PC (DMPC) | Avanti Polar Lipids | Cat#850345; CAS: 18194-24-6 |
| 1,2-distearoyl-sn-glycero-3-phosphoethanolamine (DSPE) | Avanti Polar Lipids | Cat#850715; CAS: 26853-31-6 |
| 1-(4-Isothiocyanatophenyl)-3-[6,17-dihyroxy-7,10,18,21-tetraoxo- 27-[N-acetylhydroxyla-mino) 6,11,17,22-tetraazaheptaeicosane] thiourea (DFOp-NCS) | Macrocyclics | N/A |
| 1,1-dioctadecyl-3,3,3,3-tetramethylindotricarbocyanine iodide (DiR) | AAT Bioquest | Cat#22070; CAS: 100068-60-8 |
| 4’,6-diamidino-2-phenylindole dihydrochloride | Sigma-Aldrich | Cat#D8417; CAS: 28718-90-3 |
| Acetylated LDL | Invitrogen | Cat#L35354 |
| Ammonium chloride | Sigma-Aldrich | Cat#A9434; CAS: 12125-02-9 |
| Bovine serum albumin | VWR | Cat#A7906 |
| Cell trace violet | Thermo Scientific | Cat#C34557 |
| Deoxyribonuclease I | Sigma-Aldrich | Cat#DN25, CAS: 9003-98-9 |
| DRAQ5 | Invitrogen | Cat#65-0880-96 |
| Ethylenediaminetetraacetic acid disodium (EDTA) | Sigma-Aldrich | Cat#E1644; CAS: 6381-92-6 |
| Fetal bovine serum | Sigma-Aldrich | Cat#F7524-1654682 |

| | | |
|---|---|---|
| Ficoll-Paque | GE Healthcare | N/A |
| Gentamicin | Thermo Scientific | Cat#15710064 |
| Gentamicin | Centrafarm | N/A |
| GlutaMAX™ | Thermo Scientific | Cat#35050061 |
| HDL from human plasma | Bioresource Technology | Cat#H3025 |
| Hyaluronidase type I-S | Sigma-Aldrich | Cat#H3506 |
| Isofluorane | Ecuphar | 100% v/v |
| Lipopolysaccharide (LPS) from *Escherichia coli* O55:B5 | Sigma-Aldrich | Cat#L2880 |
| Liberase TH | Sigma-Aldrich | Cat#5401151001 |
| Pam3Cys-Ser-(Lys)4 (P3C) | EMC microcollections GmbH | Cat#L2000 |
| Pancoll | PAN-Biotech | Cat#P04-601000 |
| Paraformaldehyde | VWR | Cat#43368.9M |
| Percoll | Sigma-Aldrich | Cat# P1644 |
| Penicillin / streptomycin | Sigma-Aldrich | Cat#P4333 |
| Phosphatidylserine | Avanti Polar Lipids | Cat#840032, CAS: 840032 |
| RPMI 1640 | Gibco | Cat#21875-034 |
| RPMI 1640 | Invitrogen | Cat#11875093 |
| RPMI 1640 Dutch modified medium | Thermo Scientific | Cat#22409031 |
| Sodium pyruvate | Thermo Scientific | Cat#11360070 |
| Streptavidin-phycoerythrin | BioLegend | Cat#405204 |
| Zap-Oglobin II | Beckman Coulter | Cat#7546138 |
| Zombie yellow | Biolegend | Cat#423104 |
| [$^{3}$H]-cholesterol | PerkinElmer | Cat# NET139250UC |
| [$^{18}$F]-2-deoxy-2-fluoro-D-glucose ($^{18}$F-FDG) | Curium Pharma | N/A |
| Zirconium ($^{89}$Zr) | Perkin Elmer | N/A |
| Western Diet 0.21% cholesterol | Ssniff | Cat#E15721-347 EF TD88137 |
| **Commercial assays** | | |
| BCA test | Thermo Scientific | Cat#23225 |
| Chromium Next GEM Chip G, 10x Genomics | Illumina | N/A |
| Chromium Next GEM Single Cell 3' Kit v3.1, 10x Genomics | Illumina | N/A |
| ELISA kit: DuoSet ELISA for IL-6 and TNF-α | R&D Systems | Cat#DY210 |
| Human IL-1B ELISA Kit | RD Systems | DY201 |
| Human IL-1Ra ELISA Kit | RD Systems | DRA00B |
| Human IL-6 ELISA Kit | Sanquin | M9316 |
| Human IL-18 ELISA Kit | Simple Plex | SPCKB-PS-000501 |
| Human IL-18BP ELISA Kit | R&D Duoset | DBP180 |
| Human AAT ELISA Kit | R&D Systems | DY1268 |
| Human Adiponectin ELISA Kit | R&D Systems | DY1065 |
| Human Leptin ELISA Kit | R&D Systems | DY398 |
| Human Resistin ELISA Kit | R&D Systems | DY1359 |
| Human TNF-α ELISA Kit | RD Systems | DY210 |
| Plasma IL-6 ELISA Kit | Simple Plex | SPCKB-PS-000190 |
| Plasma hsCRP ELISA Kit | R&D Systems | DY1707 |
| Plasma VEGF ELISA Kit | Simple Plex | SPCKB-PS-000330 |
| Fluorometric assay kit | Sigma-Aldrich | Cat#MAK371 |
| High sensitivity DNA chip | Agilent Technologies | N/A |
| P3 flow cell | NextSeq 2000 | N/A |
| **Experimental models** | | |
| Mice: Apoe$^{-/-}$ (females) | Charles River Laboratories | B6.129P2-*Apoe*$^{tm1Unc}$/J |

| | | |
|---|---|---|
| Mice: wild type (females) | Charles River Laboratories | C57BL/6J |
| Mice: SJL (males) | Charles Rives Laboratories | BL/6.SJLC57 (CD45.1) |
| Mice: DsRed (males) | Vintersten et al. 2004[20] | DsRed (CD45.2) |
| **Software and algorithms** | | |
| 10X CellRanger pipeline (v 6.1.1) | 10x Genomics | |
| BD FACSDiva™ (v 6.2) | BD Biosciences | |
| BD FACSDiva™ (v 6.1.3) | BD Biosciences | |
| CellBender software (v 0.2.2) | Broad Institute | |
| Cellranger mkfastq pipeline | 10x Genomics | |
| FlowJo v 10.9 | BD Biosciences | |
| GraphPad Prism 8.0. | GraphPad | |
| ImageJ | NIH | |
| LipidXplorer suite | LIFS | |
| NDPview 2 | Hamamatsu | |
| R packages | | |
| Scater packages | | |
| ScDdIFinder (v 1.12.0) | | |
| Seurat packages | | |
| **Other** | | |
| Automatic gamma counter | PerkinElmer | Wizard 1470 |
| AxioScan | Carl Zeiss | Z1 |
| BD LSRFortessa™ cytometer | BD Biosciences | LSRFortessa |
| Bioanalyzer | Agilent Technologies | 2100 |
| Cell counter | Thermo Scientific | Countess 3 |
| Cell counter (for immune cells) | Sysmex | XE-5000 |
| Column for HPLC PD10 | Cytiva | 17085101,00 |
| Column for HPLC Superose 6 10/300 GL | Cytiva | 20-0915-96 |
| Dose calibrator | Biodex | AtomLab500 |
| Dynamic light scattering | Malvern | Zetasizer Nano ZS90 |
| FACS Aria II cell sorter | BD Biosciences | Aria II |
| Haematology analyser | Horiba Medical | ABX Pentra 80 v1.12.0 |
| HPLC | Jasco | 1200 |
| Integrated chemistry system | Siemens | Dimension RxL Max |
| Q Exactive™ Plus Hybrid Quadrupole-Orbitrap™ Mass Spectrometer | Thermo Scientific | IQLAAEGAAPFALGMBDK |
| Multi-Purpose Scintillation Counter | Beckman Counter | LS 6500 |
| Photodiode array detector | Jasco | SPD-M10AVP |
| Polystyrene flat-bottom plates | Corning | CLS3599 |
| Pumps for HPLC | Jasco | LC-10AT |
| Qubit fluorometer | Thermo Scientific | |
| Scale | Sartorius | 37070 |
| Strainer 40 µm | vWR | 734-2760 |
| Strainer 70 µm | vWR | 734-2761 |
| Thermal cycler | Agilent Technologies | SureCycler 8800 |
| Ultrasonic homogenizer | Bandelin | Soloplus HD 2070 |

**Table S1.** Detected lipid classes in the 300OB cohort and average plasma concentration.

| Lipid classes | Mean concentration ± SD [µmol/L] |
|---|---|
| Cholesterol esters | 11926 ± 3934 |
| Ceramides | 11.9 ± 3.6 |
| Diglyceride | 58.2 ± 33.5 |
| DiHexosylceramide | 13.2 ± 4.4 |
| Hexosylceramides | 14.3 ± 4.8 |
| Lysophosphatidylcholines | 285 ± 67 |
| Ether-Linked lysophosphatidylcholines | 1.8 ± 0.8 |
| Lysophosphatidylethanolamine | 8.8 ± 2.5 |
| Monoglyceride | 39.9 ± 69.8 |
| Phosphatidylcholines | 5860 ± 2285 |
| Ether-Linked Phosphatidylcholines | 285 ± 118 |
| Phosphatidylethanolamine | 41.8 ± 19.8 |
| Ether-Linked Phosphatidylethanolamine | 9 ± 3.7 |
| Phosphatidylglycerol | 8.6 ± 9.8 |
| Phosphatidylserine | 68.4 ± 66.3 |
| Sphingomyelin | 431 ± 103 |
| Triglyceride | 1941 ± 1171 |

**Table S2.** Radioactivity distribution in selected tissues measured by gamma counting at 24 hours p.i. of $^{89}$Zr-labeled PS@rHDL in $WD_{6w}$ and $WD_{14w}$ mice (n = 4-5 per group).

| Tissue | %ID/g $WD_{6w}$ | %ID/g $WD_{14w}$ |
|---|---|---|
| bone marrow | 9.7 ± 4.0 | 18.0 ± 5.0 |
| liver | 9.6 ± 1.3 | 11.6 ± 3.3 |
| spleen | 5.9 ± 1.2 | 7.7 ± 1.0 |
| kidney | 6.9 ± 0.9 | 7.9 ± 0.4 |
| heart | 0.48 ± 0.06 | 0.57 ± 0.12 |
| aorta | 0.69 ± 0.18 | 1.4 ± 0.3 |
| lungs | 0.34 ± 0.14 | 0.6 ± 0.3 |
| muscle | 0.39 ± 0.15 | 1.2 ± 0.4 |
| skin | 0.47 ± 0.13 | 0.85 ± 0.65 |
| adipose tissue | 0.28 ± 0.24 | 0.19 ± 0.10 |
| mineral bone | 1.8 ± 0.2 | 3.9 ± 2.5 |
| brain | 0.02 ± 0.01 | 0.04 ± 0.01 |
| blood | 0.67 ± 0.21 | 1.3 ± 0.3 |
| plasma | 0.86 ± 0.24 | 1.1 ± 0.2 |
| blood cells | 0.30 ± 0.13 | 0.44 ± 0.19 |

**A**

| | rHDL | PS@rHDL |
|---|---|---|
| Size [d.nm] | 9.8 ± 1.6 | 10.2 ± 1.4 |
| Dispersity | 0.20 ± 0.05 | 0.25 ± 0.07 |
| Z-pot. [mV] | -8.5 ± 1.9 | -20.9 ± 8.0 |

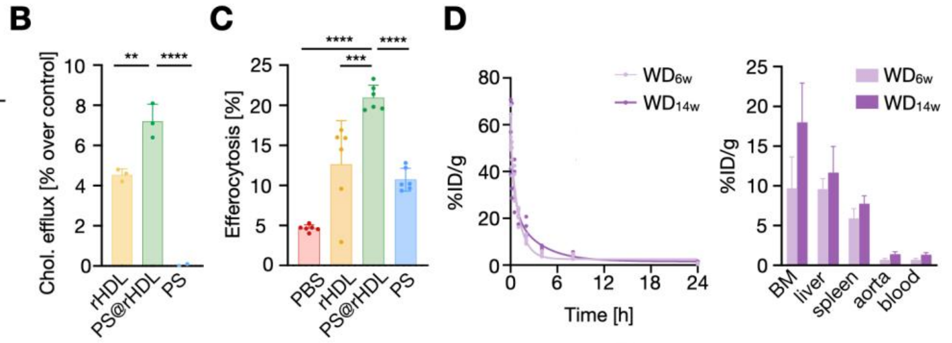


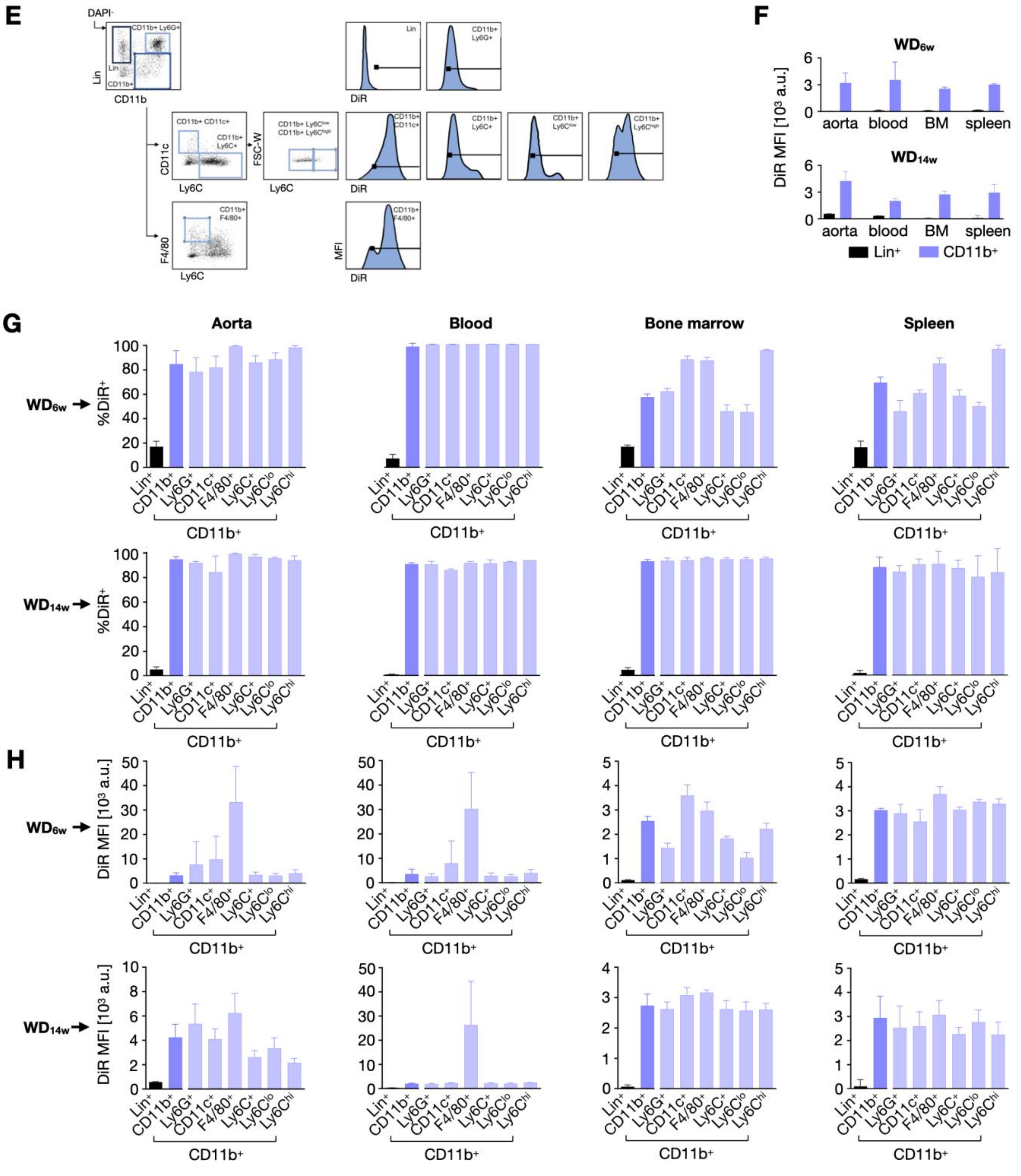

**Figure S1. PS@rHDL characterization. A.** Size, dispersity and z-potential of rHDL and PS@rHDL. **B.** Percentage of cholesterol efflux of BMDMs treated with rHDL, PS@rHDL or PS. **C.** Percentage of efferocytosis of BMDMs treated with rHDL, PS@rHDL or PS. **D.** *Ex vivo* radioactivity characterization of PS@rHDL. Blood time-activity curves for intravenously infused $^{89}$Zr-labeled PS@rHDL in $WD_{6w}$ and $WD_{14w}$ mice (left, n = 4-5 per group). The weighted half-lives for $WD_{6w}$ and $WD_{14w}$ mice were determined to be 37 and 72 min, respectively. Biodistribution in tissues of interest measured by gamma counting at 24 hours p.i. of $^{89}$Zr-labeled PS-rHDL in $WD_{6w}$ and $WD_{14w}$ mice (right, n = 4-5 per group). **E**. Gating procedure employed to determine cell targeting specificity in the aorta, blood, bone marrow and spleen of $WD_{6w}$ and $WD_{14w}$ mice using flow cytometry and fluorescent DiR-labeled PS@rHDL. **F**. Quantification of DiR accumulation in different cell types in the aorta, blood, bone marrow and spleen of $WD_{6w}$ and $WD_{14w}$ mice (n = 4 per group). In all tissues, PS@rHDL showed selectivity for myeloid cells ($CD11b^+$), measured as both percentage of DiR-positive cells (**G**) and as median fluorescent intensity (MFI, **H**). In the aorta, macrophages ($CD11b^+F4/80^+$) were preferentially targeted over the other populations, especially in the $WD_{6w}$ group. Data are presented as mean ± SD, and P values were calculated using one-way ANOVA followed by multiple comparison using Dunnett's test. ** $P < 0.01$, *** $P < 0.001$ and **** $P < 0.0001$. $WD_{6w}$: *Apoe*$^{-/-}$ mice fed a WD for 6 weeks; $WD_{14w}$: *Apoe*$^{-/-}$ mice fed a WD for 14 weeks.

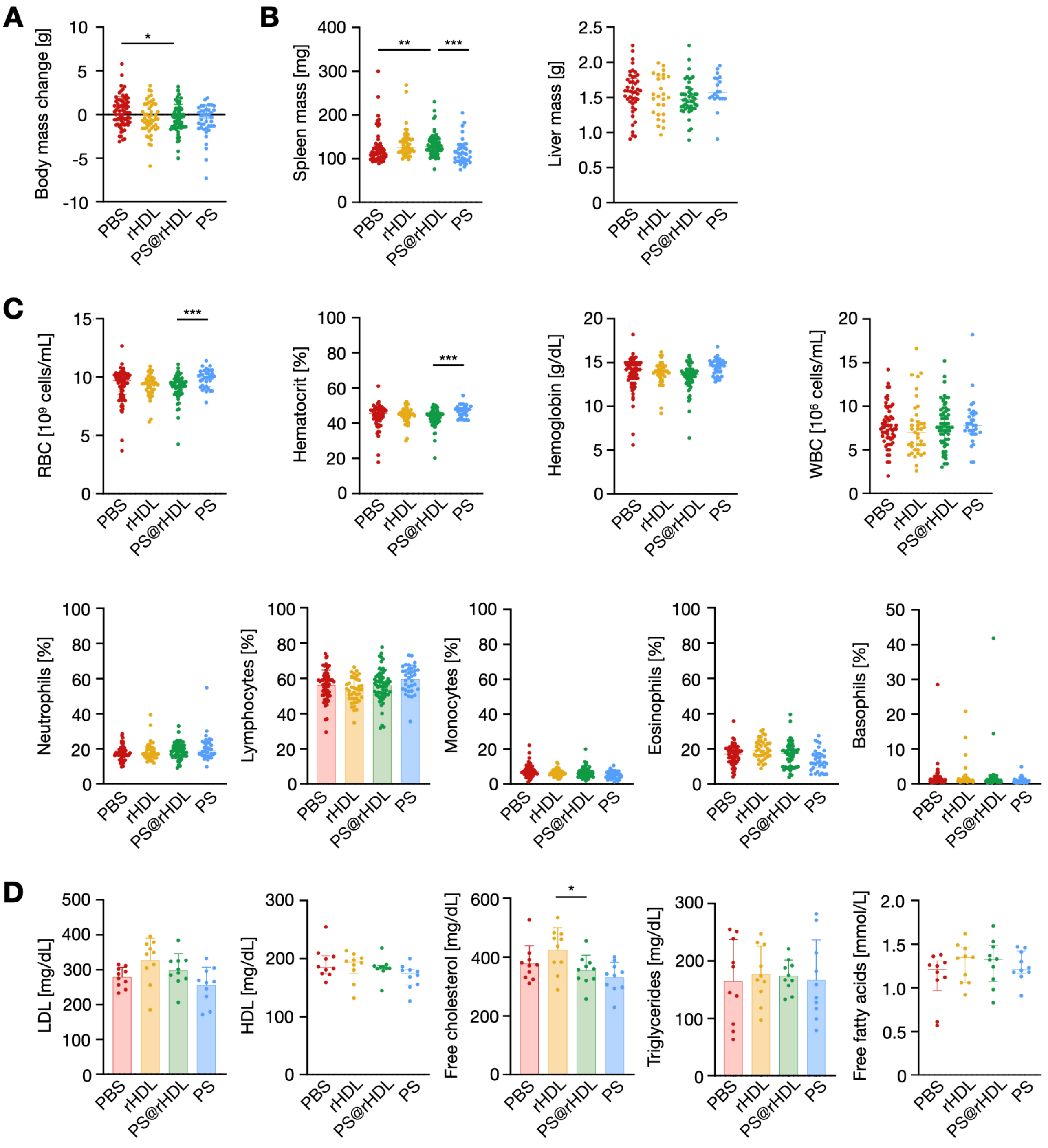


**Figure S2. PS@rHDL treatment is well tolerated [$WD_{6w}$].** Assessment of PS@rHDL's potential adverse effects in $WD_{6w}$ mice. **A.** Body mass change (n ≥ 40 per group). **B.** Spleen (left, n ≥ 40 per group) and liver (right, n ≥ 25 per group) mass after treatment. **C.** Complete blood count hematological analysis after treatment showing red blood cell (RBC) numbers, hematocrit, hemoglobin, and white blood cell (WBC) numbers, as well as neutrophil, lymphocyte, monocyte, eosinophil and basophil percentages (n ≥ 35 per group). **D.** Lipid profile after treatment showing LDL, HDL, free cholesterol, total cholesterol, triglyceride and free fatty acid plasma concentrations (n = 10 per group). Normally distributed data are presented as mean ± SD, and P values were calculated using one-way ANOVA, followed by multiple comparisons using Dunnett's test. Otherwise, data are presented as median and IQR, with P values calculated using Kruskal-Wallis test, followed by multiple comparison using Dunn's test. * $P < 0.05$, ** $P < 0.01$ and *** $P < 0.001$. RBC: red blood cell; WBC: white blood cell; LDL: low-density lipoprotein; HDL: high-density lipoprotein.

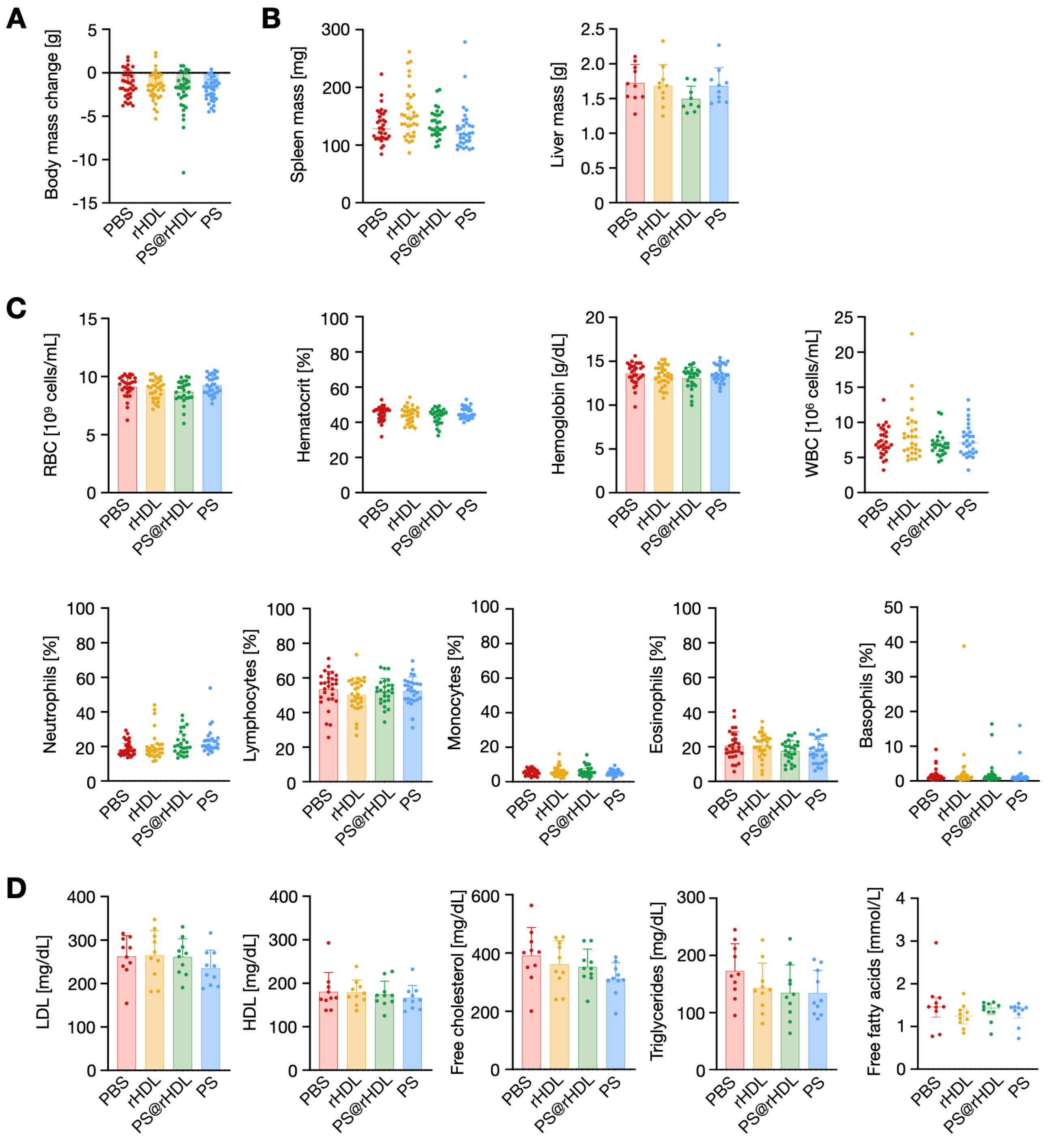


**Figure S3. PS@rHDL treatment is well tolerated [$WD_{14w}$].** Assessment of PS@rHDL's potential adverse effects in $WD_{14w}$ mice. **A.** Body mass change (n = 35 per group). **B.** Spleen (left, n = 35 per group) and liver (right, n = 10 per group) mass after treatment. **C.** Complete blood count hematological analysis after treatment showing red blood cell (RBC) numbers, hematocrit, hemoglobin, and (WBC) numbers, as well as neutrophil, lymphocyte, monocyte, eosinophil and basophil percentages (n = 30 per group). **D.** Lipid profile after treatment showing LDL, HDL, free cholesterol, total cholesterol, triglyceride and free fatty acid plasma concentrations (n = 10 per group). Normally distributed data are presented as mean ± SD, and P values were calculated using one-way ANOVA, followed by multiple comparisons using Dunnett's test. Otherwise, data are presented as median and IQR, with P values calculated using Kruskal-Wallis test, followed by multiple comparison using Dunn's test. RBC: red blood cell; WBC: white blood cell; LDL: low-density lipoprotein; HDL: high-density lipoprotein.

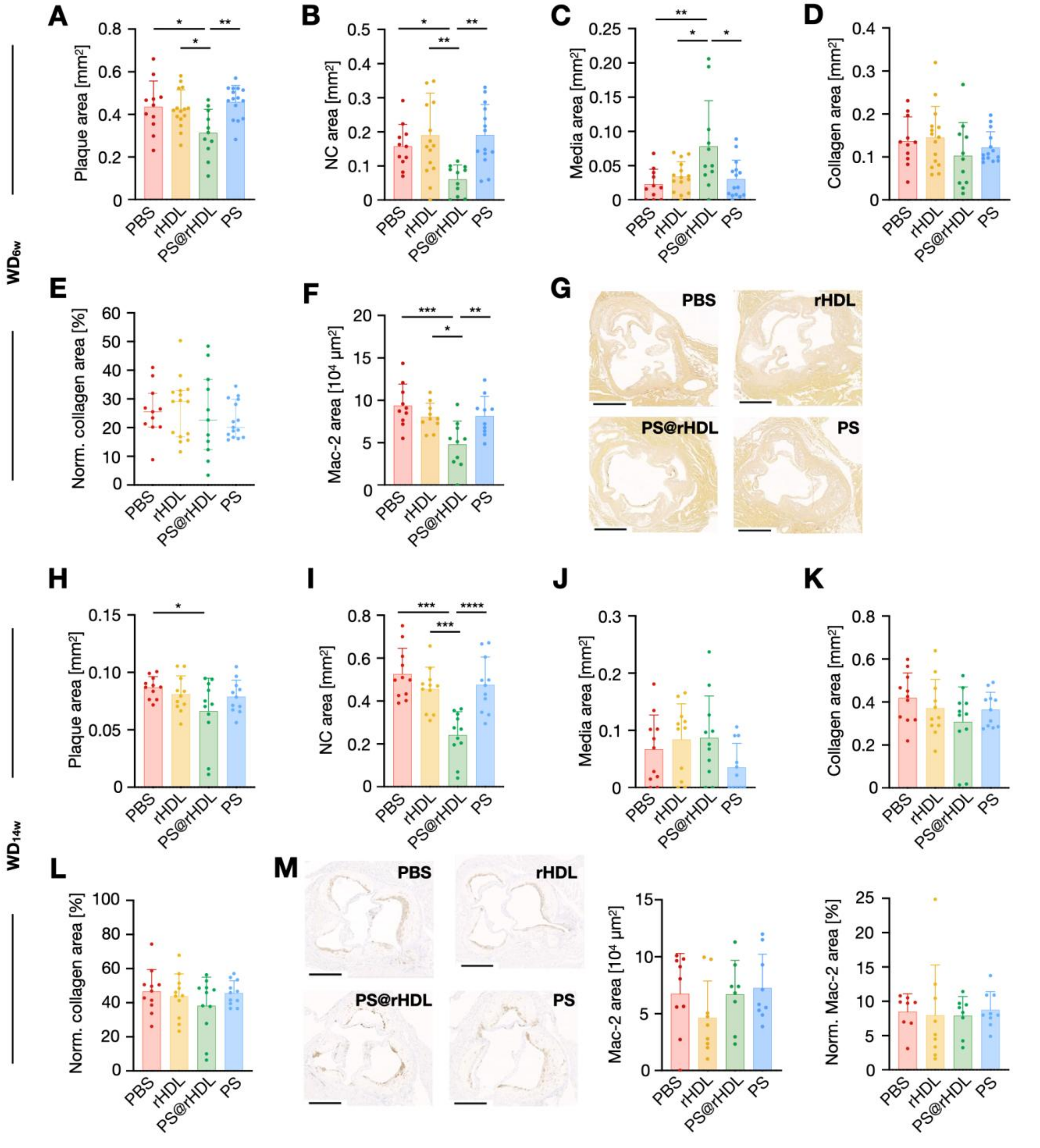


**Figure S4. PS@rHDL reduces necrotic core and plaque size in atherosclerosis [II]. A, B, C.** Quantitative analysis of plaque (**A**), necrotic core (**B**), and tunica media (**C**) areas in aortic roots from treated WD$_{6w}$ mice (n = 10-15 per group). **D, E.** Collagen area (**D**) and normalized collagen area (**E**) in aortic roots from treated WD$_{6w}$ mice (n = 10-15 per group). **F.** Quantitative analysis of Mac-2$^+$ area in aortic roots from treated WD$_{6w}$ mice (n = 10 per group). **G.** Representative images of Alizarin red-stained sections from atherosclerotic lesions in aortic roots from treated WD$_{6w}$ mice (n = 10 per group). Scale bar = 500 µm. **H, I, J.** Quantitative analysis of plaque area (**H**), necrotic core (**I**), and tunica media (**J**) areas in aortic roots from treated WD$_{14w}$ mice (n = 11 per group). **K, L.** Collagen area (**K**) and normalized collagen area (**L**) in aortic roots from treated WD$_{14w}$ mice (n = 11 per group). **M.** Representative Mac-2 immunohistochemistry images and quantitative analysis of Mac-2$^+$ area and normalized Mac-2$^+$ area of atherosclerotic lesions in aortic roots from treated WD$_{14w}$ (n = 8-9 per group). Scale bar = 500 µm. Normally distributed data are presented as mean ± SD, and P values were calculated using one-way ANOVA, followed by multiple comparisons using Dunnett's test. Otherwise, data are presented as median and IQR, with P values calculated using Kruskal-Wallis test, followed by multiple comparison using Dunn's test. * $P < 0.05$, ** $P < 0.01$, *** $P < 0.001$ and **** $P < 0.0001$. WD$_{6w}$: *Apoe*$^{-/-}$ mice fed a WD for 6 weeks; WD$_{14w}$: *Apoe*$^{-/-}$ mice fed a WD for 14 weeks; NC: necrotic core.

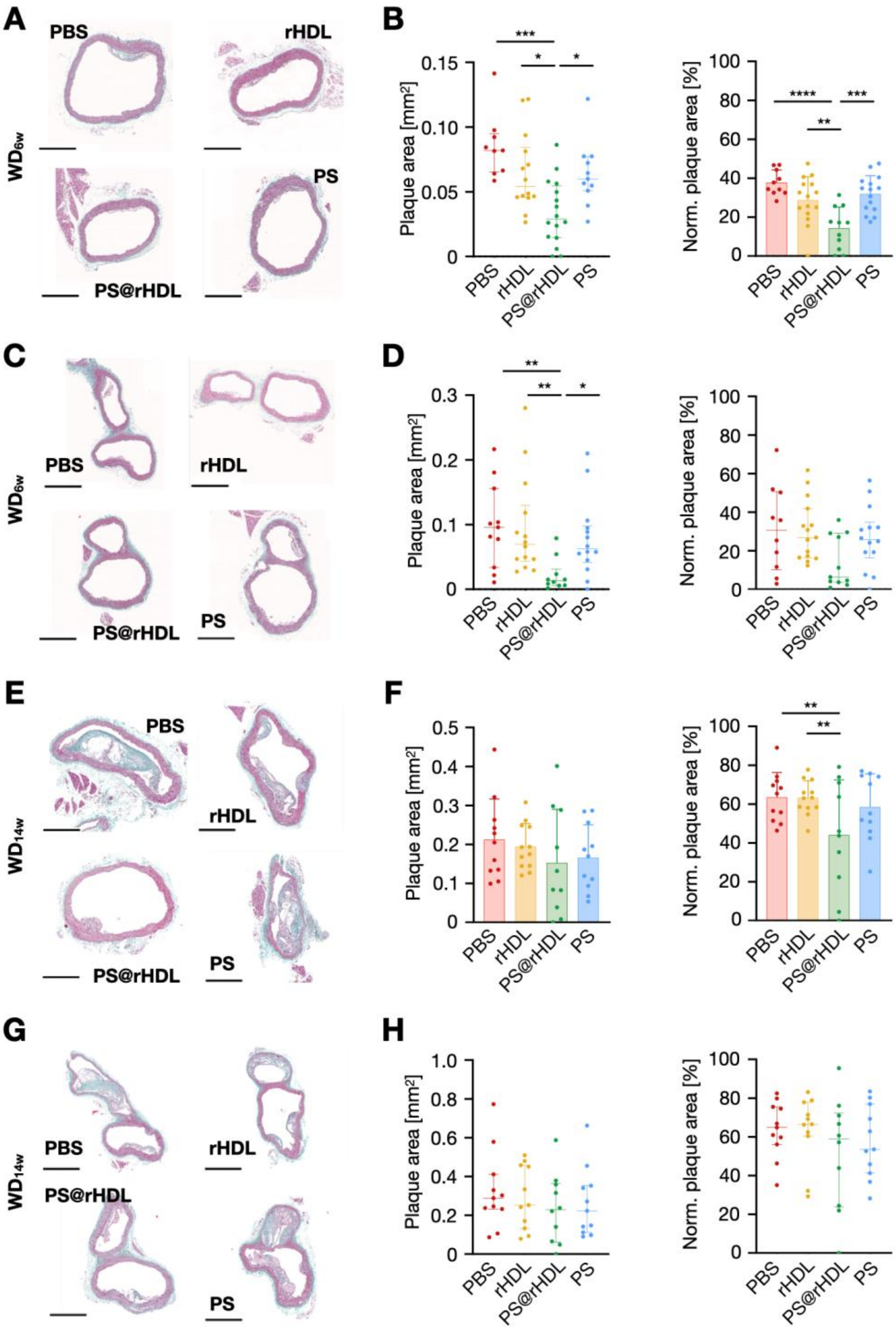


**Figure S5. PS@rHDL reduces necrotic core and plaque size in atherosclerosis [III]. A, B.** Representative images of atherosclerotic lesions in the ascending aorta stained with Masson's Trichrome (**A**), and corresponding quantitative analysis of plaque area (**B**, left) and normalized plaque area (**B**, right) from $WD_{6w}$ mice (n = 10-15 per group). Scale bar = 500 µm. **C, D.** Representative images of atherosclerotic lesions in the aortic-brachiocephalic artery bifurcation stained with Masson's Trichrome (**C**), and corresponding quantitative analysis of plaque area (**D**, left) and normalized plaque area (**D**, right) from $WD_{6w}$ mice (n = 10-15 per group). Scale bar = 500 µm. **E, F.** Representative images of atherosclerotic lesions in the ascending aorta stained with Masson's Trichrome (**E**), and corresponding quantitative analysis of plaque area (**F**, left) and normalized plaque area (**F**, right) from $WD_{14w}$ mice (n = 11 per group). Scale bar = 500 µm. **G, H.** Representative images of atherosclerotic lesions in the aortic-brachiocephalic artery bifurcation stained with Masson's Trichrome (**G**), and corresponding quantitative analysis of plaque area (**H**, left) and normalized plaque area (**H**, right) from $WD_{14w}$ mice (n = 11 per group). Scale bar = 500 µm. Normally distributed data are presented as mean ± SD, and P values were calculated using one-way ANOVA, followed by multiple comparisons using Dunnett's test. Otherwise, data are presented as median and IQR, with P values calculated using Kruskal-Wallis test, followed by multiple comparison using Dunn's test. * $P < 0.05$, ** $P < 0.01$, *** $P < 0.001$ and **** $P < 0.0001$. $WD_{6w}$: *Apoe*$^{-/-}$ mice fed a WD for 6 weeks; $WD_{14w}$: *Apoe*$^{-/-}$ mice fed a WD for 14 weeks.

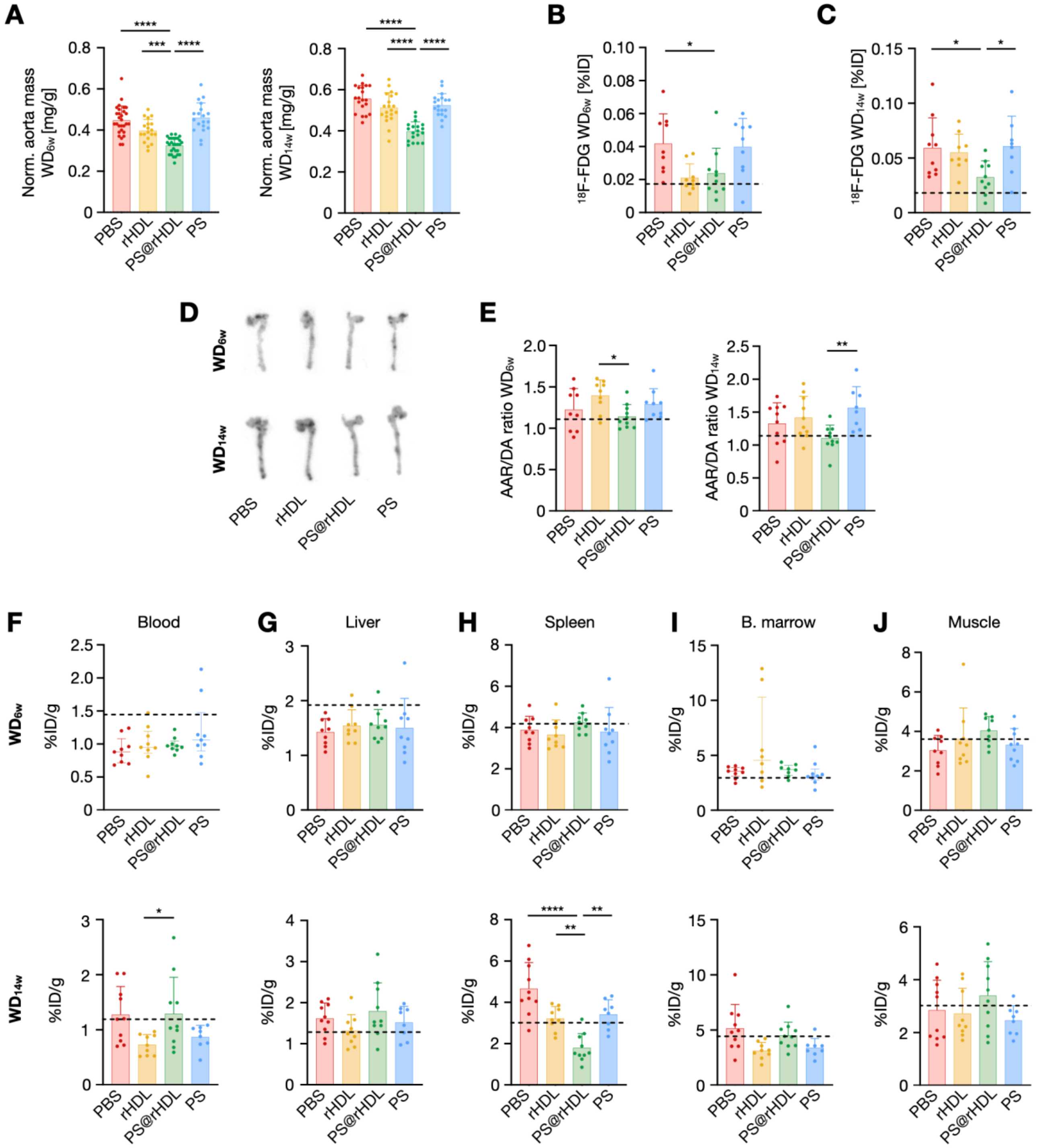


**Figure S6. PS@rHDL normalizes vessel wall metabolism and architecture [II]. A.** Aortic mass normalized to body weight from treated WD$_{6w}$ (left, n = 20-30 per group) and WD$_{14w}$ (right, n = 20 per group) mice. **B, C.** $^{18}$F-FDG uptake in aortas from treated WD$_{6w}$ (**B**) and WD$_{14w}$ mice (**C**) expressed as percentage of injected dose (%ID). **D.** Representative autoradiographs of aortas from treated WD$_{6w}$ (top) and WD$_{14w}$ mice (bottom) showing regional distribution of $^{18}$F-FDG uptake. **E.** Aortic arch & root (AAR)-to-descending aorta (DA) $^{18}$F-FDG uptake ratios as measured from autoradiographs of aortas from treated WD$_{6w}$ (left) and WD$_{14w}$ mice (right). **F-J.** $^{18}$F-FDG uptake in blood (**F**), liver (**G**), spleen (**H**), bone marrow (**I**) and muscle (**J**) from WD$_{6w}$ (top row) and WD$_{14w}$ mice (bottom row) after treatment (n = 10 per group). Dashed black lines represent average values from age-matched wild type C57BL/6J mice. Normally distributed data are presented as mean ± SD, and P values were calculated using one-way ANOVA, followed by multiple comparisons using Dunnett's test. Otherwise, data are presented as median and IQR, with P values calculated using Kruskal-Wallis test, followed by multiple comparison using Dunn's test. * $P < 0.05$, ** $P < 0.01$, *** $P < 0.001$ and **** $P < 0.0001$. WD$_{6w}$: *Apoe*$^{-/-}$ mice fed a WD for 6 weeks; WD$_{14w}$: *Apoe*$^{-/-}$ mice fed a WD for 14 weeks.

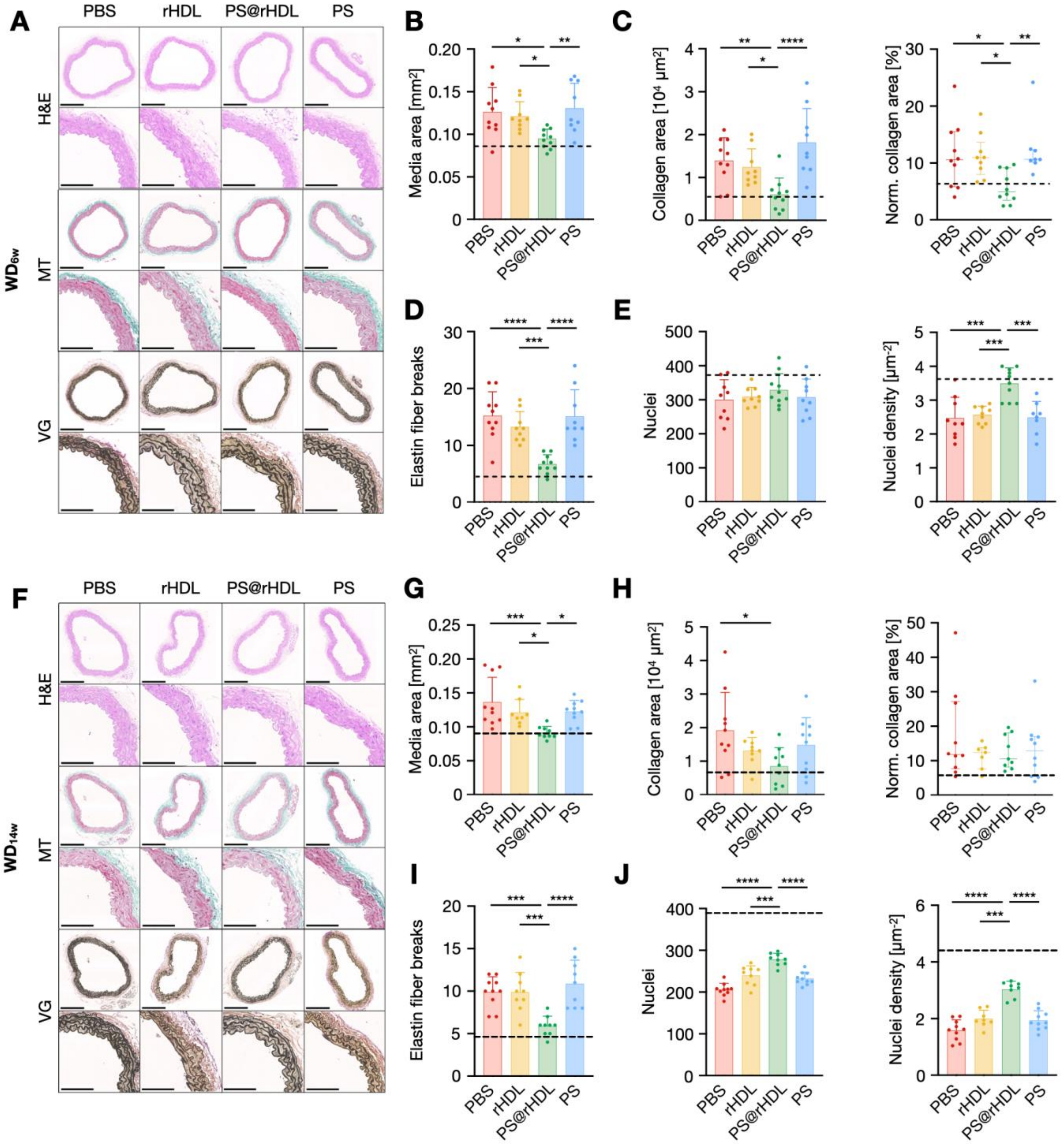


**Figure S7. PS@rHDL normalizes vessel wall metabolism and architecture [III]. A.** Representative images showing hematoxylin & eosin (H&E), Masson's Trichrome (MT) and Van Gieson (VG) stainings of the vessel wall from the lower section of the descending thoracic aorta from treated $WD_{6w}$ mice (scale bar = 250 μm) and their respective magnifications (scale bar = 100 μm) after treatment. **B, C, D, E.** Quantitative analysis of tunica media area (**B**), collagen area (**C**, left), normalized collagen area (**C**, right), number of elastin fiber breaks (**D**), number of nuclei (**E**, left) and nuclei density (**E**, right) in the lower section of the descending thoracic aorta from treated $WD_{6w}$ mice (n = 10 per group). **F.** Representative images showing hematoxylin & eosin (H&E), Masson's Trichrome (MT) and Van Gieson (VG) stainings of the vessel wall from the lower section of the descending thoracic aorta from treated $WD_{14w}$ mice (scale bar = 250 μm) and their respective magnifications (scale bar = 100 μm). **G, H, I, J.** Quantitative analysis of tunica media area (**G**), collagen area (**H**, left), normalized collagen area (**H**, right), number of elastin fiber breaks (**I**), number of nuclei (**J**, left) and nuclei density (**J**, right) in the lower section of the descending thoracic aorta from treated $WD_{14w}$ mice (n = 10 per group). Dashed black lines represent average values from age-matched wild type C57BL/6J mice. Normally distributed data are presented as mean ± SD, and P values were calculated using one-way ANOVA, followed by multiple comparisons using Dunnett's test. Otherwise, data are presented as median and IQR, with P values calculated using Kruskal-Wallis test, followed by multiple comparison using Dunn's test. * $P < 0.05$, ** $P < 0.01$, *** $P < 0.001$ and **** $P < 0.0001$. $WD_{6w}$: *Apoe*$^{-/-}$ mice fed a WD for 6 weeks; $WD_{14w}$: *Apoe*$^{-/-}$ mice fed a WD for 14 weeks; H&E: hematoxylin & eosin; MT: Masson's Trichrome; VG: Van Gieson.

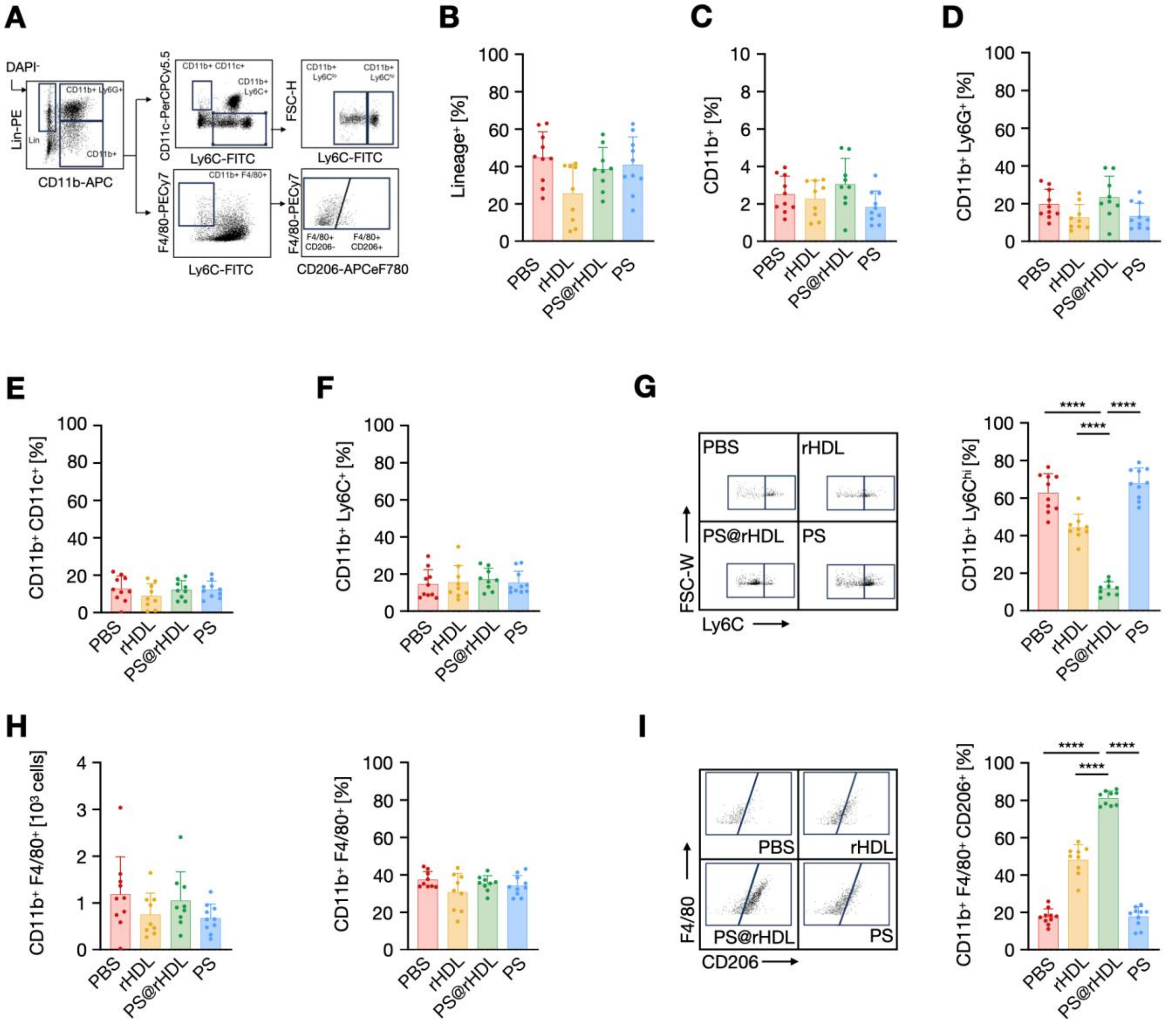


**Figure S8. PS@rHDL modulates the monocyte/macrophage compartment [Aorta $WD_{6w}$]. A.** Representative flow cytometry gating strategy to identify relevant immune cells in aorta, bone marrow, blood and spleen after treatment. **B, C, D, E, F.** Frequency of Linage$^+$ cells (including CD90.2$^+$, CD49b$^+$, NK1.1$^+$, B220$^+$ or Ter119$^+$ cells, **B**), myeloid cells (CD11b$^+$ cells, **C**), neutrophils (CD11b$^+$Ly6G$^+$ cells, **D**), dendritic cells (CD11b$^+$CD11c$^+$ cells, **E**) and monocytes (CD11b$^+$Ly6C$^+$ cells, **F**) in aortas from treated $WD_{6w}$ mice. **G.** Representative plots showing the proportion of aortic Ly6C$^{hi}$ monocytes (left) and corresponding quantification in each treatment group (right, n = 10 per group) in treated $WD_{6w}$ mice. **H.** Number (left) and frequency (right) of macrophages (identified as CD11b$^+$F4/80$^+$ cells) in aortas from treated $WD_{6w}$ mice. **I.** Representative plots showing the proportion of aortic CD206$^+$ macrophages (left) and corresponding quantification in each treatment group (right) in treated $WD_{6w}$ mice (n = 10 per group). Normally distributed data are presented as mean ± SD, and P values were calculated using one-way ANOVA, followed by multiple comparisons using Dunnett's test. **** $P < 0.0001$. $WD_{6w}$: *Apoe*$^{-/-}$ mice fed a WD for 6 weeks.

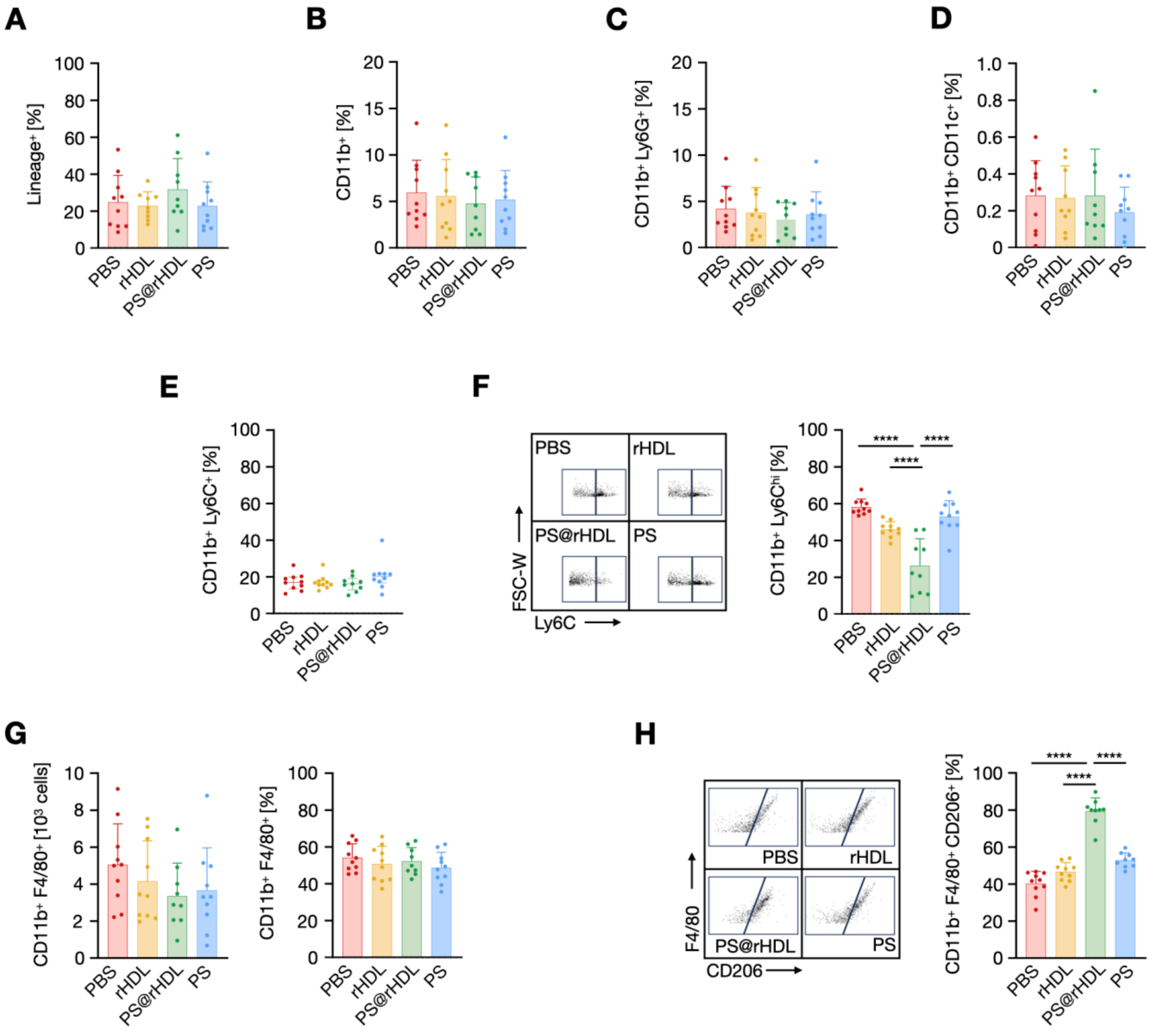


**Figure S9. PS@rHDL modulates the monocyte/macrophage compartment [Aorta $WD_{14w}$]. A, B, C, D, E.** Frequency of Linage$^+$ cells (including as CD90.2$^+$, CD49b$^+$, NK1.1$^+$, B220$^+$ or Ter119$^+$ cells, **A**), myeloid cells (CD11b$^+$ cells, **B**), neutrophils (CD11b$^+$Ly6G$^+$ cells, **C**), dendritic cells (CD11b$^+$CD11c$^+$ cells, **D**) and monocytes (CD11b$^+$Ly6C$^+$ cells, **E**) in aortas from treated $WD_{14w}$ mice (n = 10 per group). **F.** Representative plots showing the proportion of aortic Ly6C$^{hi}$ monocytes (left) and corresponding quantification in each treatment group (right) (n = 10 per group) in treated $WD_{14w}$ mice. **G.** Number (left) and frequency (right) of macrophages (identified as CD11b$^+$F4/80$^+$ cells) in aortas from treated $WD_{14w}$ mice. **H.** Representative plots showing the proportion of aortic CD206$^+$ macrophages (left) and corresponding quantification in each treatment group (right) in treated $WD_{14w}$ mice (n = 10 per group). Normally distributed data are presented as mean ± SD, and P values were calculated using one-way ANOVA, followed by multiple comparisons using Dunnett's test. Otherwise, data are presented as median and IQR, with P values calculated using Kruskal-Wallis test, followed by multiple comparison using Dunn's test **** $P < 0.0001$. $WD_{6w}$: *Apoe*$^{-/-}$ mice fed a WD for 14 weeks.

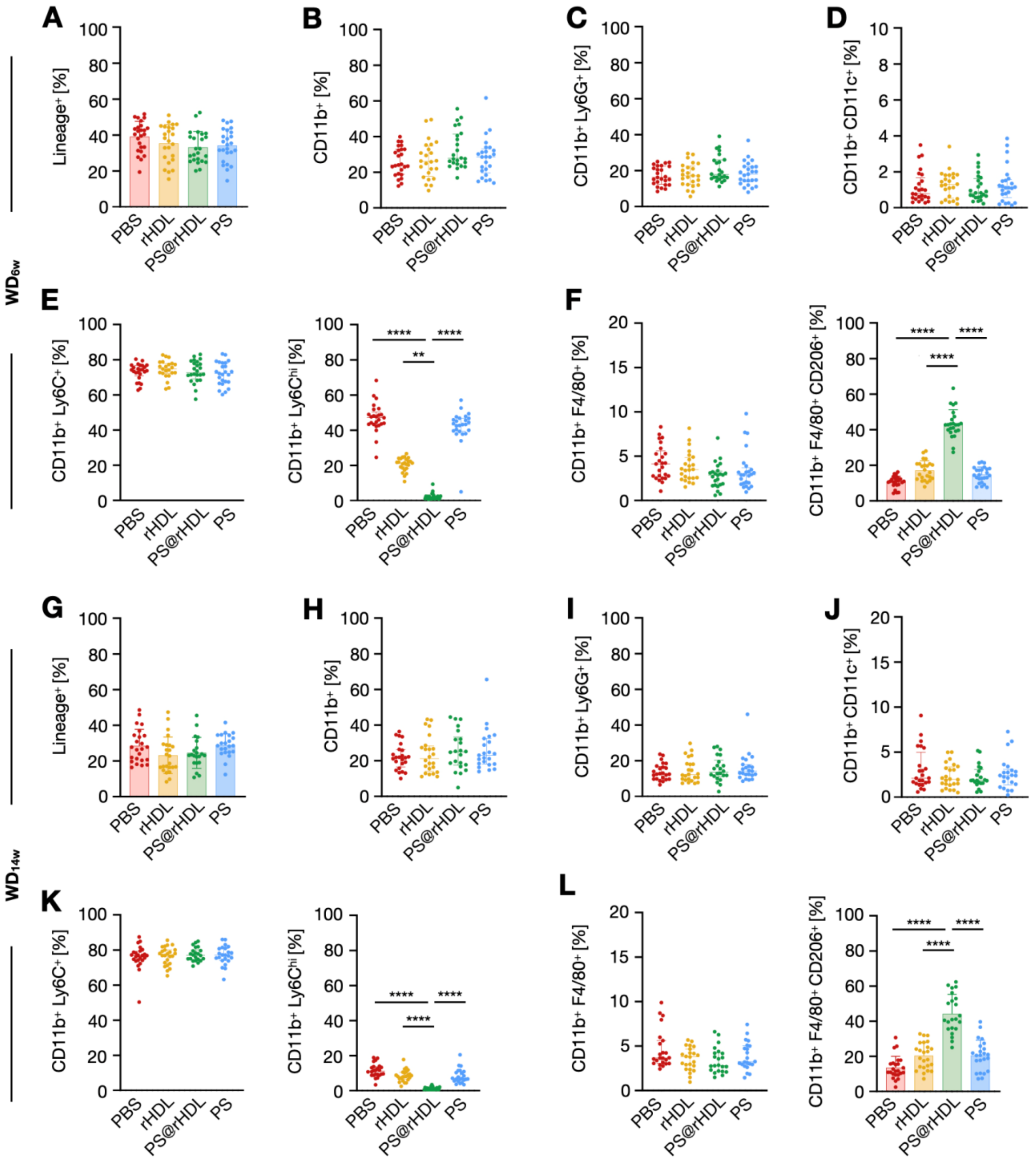


**Figure S10. PS@rHDL modulates the monocyte/macrophage compartment [Bone marrow]. A, B, C, D.** Frequency of Linage$^+$ cells (including CD90.2$^+$, CD49b$^+$, NK1.1$^+$, B220$^+$ or Ter119$^+$ cells, **A**), myeloid cells (CD11b$^+$ cells, **B**), neutrophils (CD11b$^+$Ly6G$^+$ cells,**C**) and dendritic cells (CD11b$^+$CD11c$^+$ cells, **D**) in the bone marrow from treated WD$_{6w}$. **E.** Frequency of bone marrow monocytes (CD11b$^+$Ly6C$^+$ cells, left) and Ly6C$^{hi}$ monocytes (CD11b$^+$Ly6C$^{hi}$ cells, right) in treated WD$_{6w}$. **F.** Frequency of bone marrow macrophages (CD11b$^+$F4/80$^+$ cells, left) and CD206$^+$ macrophages (CD11b$^+$F4/80$^+$CD206$^+$ cells, right) in treated WD$_{6w}$ (n = 25 per group). **G, H, I, J.** Frequency of Linage$^+$ cells (including CD90.2$^+$, CD49b$^+$, NK1.1$^+$, B220$^+$ or Ter119$^+$ cells, **G**), myeloid cells (CD11b$^+$ cells, **H**), neutrophils (CD11b$^+$Ly6G$^+$ cells, **I**) and dendritic cells (CD11b$^+$CD11c$^+$ cells, **J**) in the bone marrow from treated WD$_{14w}$. **K.** Frequency of bone marrow monocytes (CD11b$^+$Ly6C$^+$ cells, left) and Ly6C$^{hi}$ monocytes (CD11b$^+$Ly6C$^{hi}$ cells, right) in treated WD$_{14w}$. **L.** Frequency of bone marrow Frequency of bone marrow macrophages (CD11b$^+$F4/80$^+$ cells, left) and CD206$^+$ macrophages (CD11b$^+$F4/80$^+$CD206$^+$ cells, right) in treated WD$_{14w}$ mice (n = 25 per group). Normally distributed data are presented as mean ± SD, and P values were calculated using one-way ANOVA, followed by multiple comparisons using Dunnett's test. Otherwise, data are presented as median and IQR, with P values calculated using Kruskal-Wallis test, followed by multiple comparison using Dunn's test. ** $P < 0.01$ and **** $P < 0.0001$. WD$_{6w}$: *Apoe*$^{-/-}$ mice fed a WD for 6 weeks; WD$_{14w}$: *Apoe*$^{-/-}$ mice fed a WD for 14 weeks.

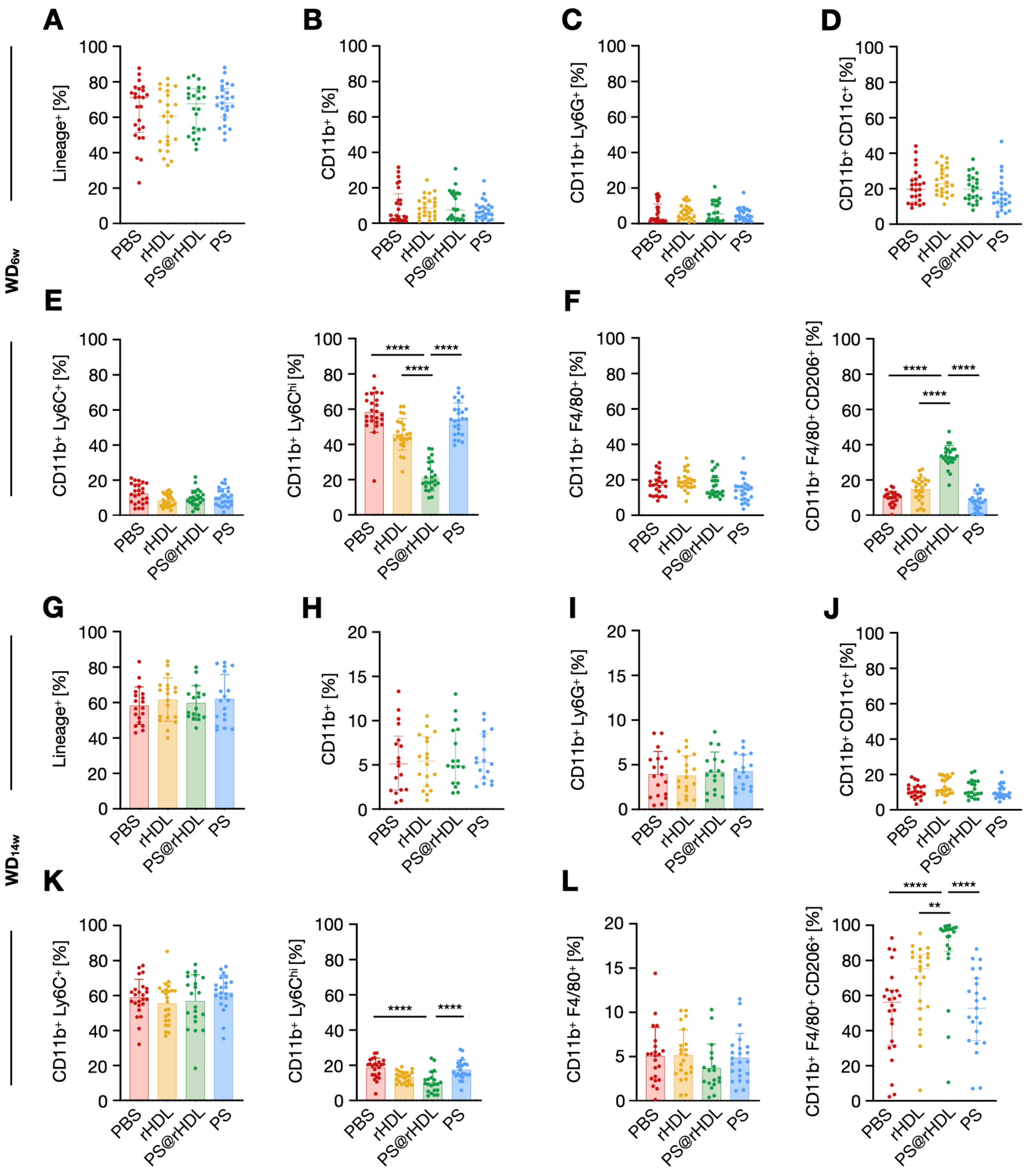


**Figure S11. PS@rHDL modulates the monocyte/macrophage compartment [Blood]. A, B, C, D.** Frequency of Linage$^+$ cells (including CD90.2$^+$, CD49b$^+$, NK1.1$^+$, B220$^+$ or Ter119$^+$ cells, **A**), myeloid cells (CD11b$^+$ cells, **B**), neutrophils (CD11b$^+$Ly6G$^+$ cells, **C**) and dendritic cells (CD11b$^+$CD11c$^+$ cells, **D**) in blood from treated WD$_{6w}$. **E.** Frequency of blood monocytes (CD11b$^+$Ly6C$^+$ cells, left) and Ly6C$^{hi}$ monocytes (CD11b$^+$Ly6C$^{hi}$ cells, right) in treated WD$_{6w}$. **F.** Frequency of blood CD11b$^+$F4/80$^+$ (left) and CD11b$^+$F4/80$^+$CD206$^+$ (right) cells in treated WD$_{6w}$ (n = 25 per group). **G, H, I, J.** Frequency of Linage$^+$ cells (including CD90.2$^+$, CD49b$^+$, NK1.1$^+$, B220$^+$ or Ter119$^+$ cells, **G**), myeloid cells (CD11b$^+$ cells, **H**), neutrophils (CD11b$^+$Ly6G$^+$ cells, **I**) and dendritic cells (CD11b$^+$CD11c$^+$ cells **J**) in blood from treated WD$_{14w}$. **K.** Frequency of blood monocytes (CD11b$^+$Ly6C$^+$ cells, left) and Ly6C$^{hi}$ monocytes (CD11b$^+$Ly6C$^{hi}$ cells, right) in treated WD$_{14w}$. **L.** Frequency of blood CD11b$^+$F4/80$^+$ (left) and CD11b$^+$F4/80$^+$CD206$^+$ (right) cells in treated WD$_{14w}$ mice (n = 25 per group). Normally distributed data are presented as mean ± SD, and P values were calculated using one-way ANOVA, followed by multiple comparisons using Dunnett's test. Otherwise, data are presented as median and IQR, with P values calculated using Kruskal-Wallis test, followed by multiple comparison using Dunn's test. ** $P < 0.01$ and **** $P < 0.0001$. WD$_{6w}$: *Apoe*$^{-/-}$ mice fed a WD for 6 weeks; WD$_{14w}$: *Apoe*$^{-/-}$ mice fed a WD for 14 weeks.

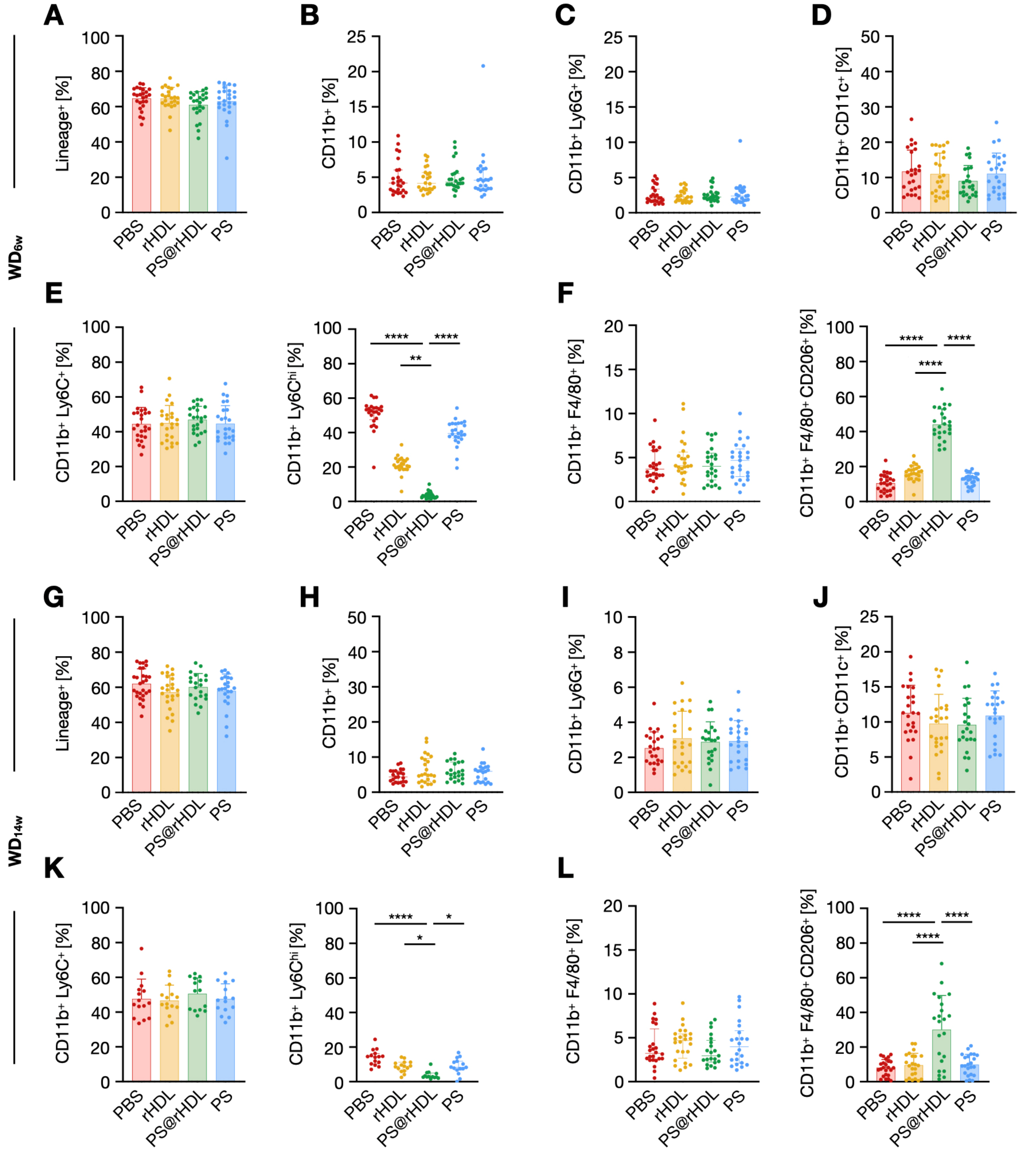


**Figure S12. PS@rHDL modulates the monocyte/macrophage compartment [Spleen]. A, B, C, D.** Frequency of Linage$^+$ cells (including CD90.2$^+$, CD49b$^+$, NK1.1$^+$, B220$^+$ or Ter119$^+$ cells, **A**), myeloid cells (CD11b$^+$ cells, **B**), neutrophils (CD11b$^+$Ly6G$^+$ cells, **C**) and dendritic cells (CD11b$^+$CD11c$^+$ cells, **D**) in the spleen from treated WD$_{6w}$. **E.** Frequency of splenic monocytes (CD11b$^+$Ly6C$^+$ cells, left) and Ly6C$^{hi}$ monocytes (CD11b$^+$Ly6C$^{hi}$ cells, right) in treated WD$_{6w}$. **F.** Frequency of splenic macrophages (CD11b$^+$F4/80$^+$ cells, left) and CD206$^+$ macrophages (CD11b$^+$F4/80$^+$CD206$^+$ cells, right) in treated WD$_{6w}$ (n = 25 per group). **G, H, I, J.** Frequency of Linage$^+$ cells (including CD90.2$^+$, CD49b$^+$, NK1.1$^+$, B220$^+$ or Ter119$^+$ cells, **G**), myeloid cells (CD11b$^+$ cells, **H**), neutrophils (CD11b$^+$Ly6G$^+$ cells, **I**) and dendritic cells (CD11b$^+$CD11c$^+$, **J**) in the spleen from treated WD$_{14w}$. **K.** Frequency of splenic monocytes (CD11b$^+$Ly6C$^+$ cells, left) and Ly6C$^{hi}$ monocytes (CD11b$^+$Ly6C$^{hi}$ cells, right) in treated WD$_{14w}$. **L.** Frequency of splenic macrophages (CD11b$^+$F4/80$^+$ cells, left) and CD206$^+$ macrophages (CD11b$^+$F4/80$^+$CD206$^+$ cells, right) in treated WD$_{14w}$ mice (n = 25 per group). Normally distributed data are presented as mean ± SD, and P values were calculated using one-way ANOVA, followed by multiple comparisons using Dunnett's test. Otherwise, data are presented as median and IQR, with P values calculated using Kruskal-Wallis test, followed by multiple comparison using Dunn's test. * $P < 0.05$, ** $P < 0.01$ and **** $P < 0.0001$. WD$_{6w}$: *Apoe$^{-/-}$* mice fed a WD for 6 weeks; WD$_{14w}$: *Apoe$^{-/-}$* mice fed a WD for 14 weeks.

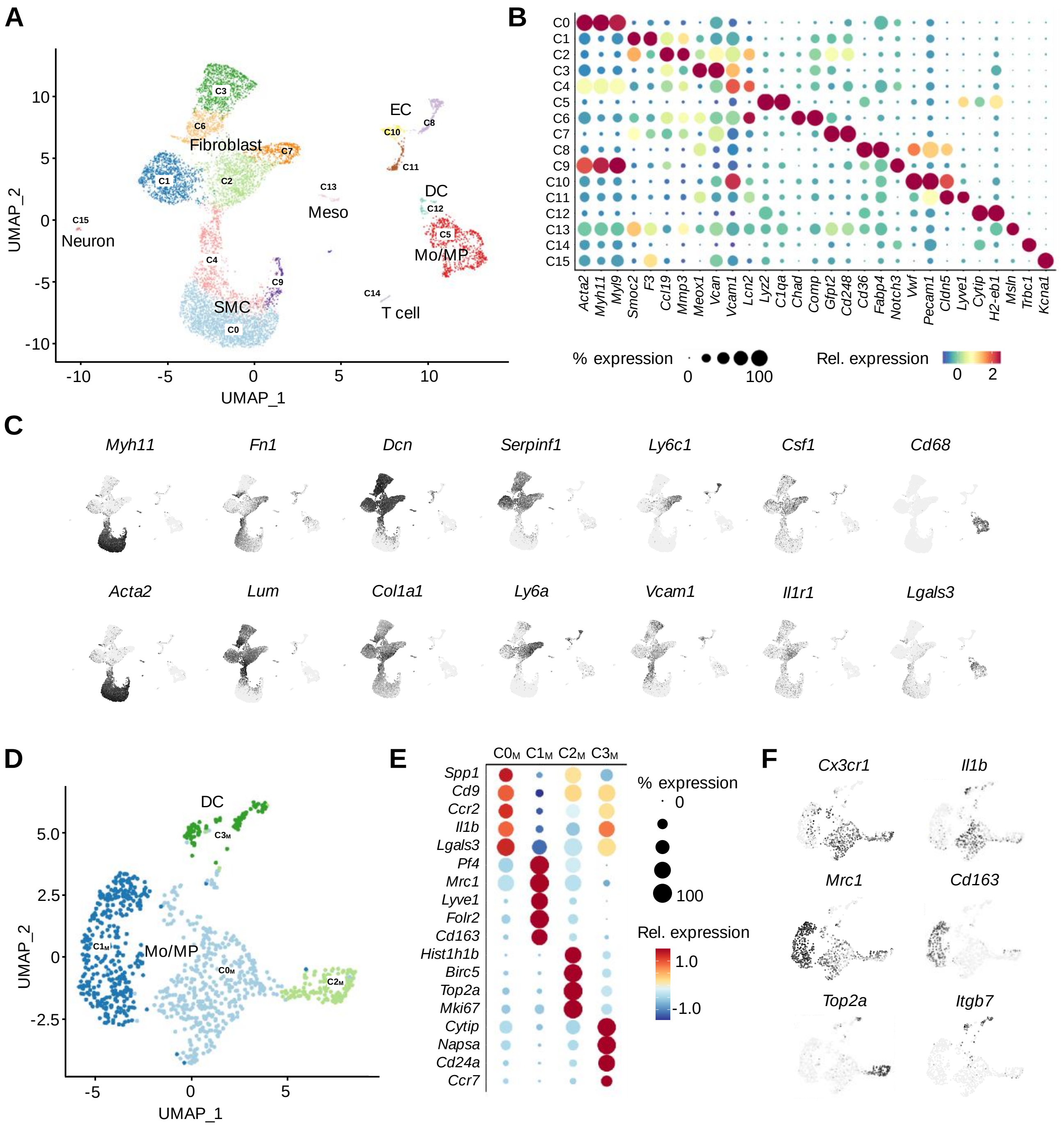


**Figure S13. Single-cell RNA sequencing maps vessel wall and plaque cellular heterogeneity. A.** Uniform manifold approximation and projection (UMAP) plot of single cells from PS@rHDL- and rHDL-treated $WD_{6w}$ mouse aortic root and arch samples displaying the 16 identified clusters. **B.** Dot plot showing a selection of representative differentially expressed genes for each cluster. **C.** UMAP plot of expression levels of selected genes used to identify and characterize relevant cellular types. **D.** UMAP plot of single cells after reclustering myeloid cell populations (C5 and C12) displaying the four resulting subclusters. **E.** Dot plot showing a selection of representative differentially expressed genes for each myeloid cell subcluster. **F.** UMAP plot of expression levels of selected genes used to identify and characterize the myeloid cell sub clusters.

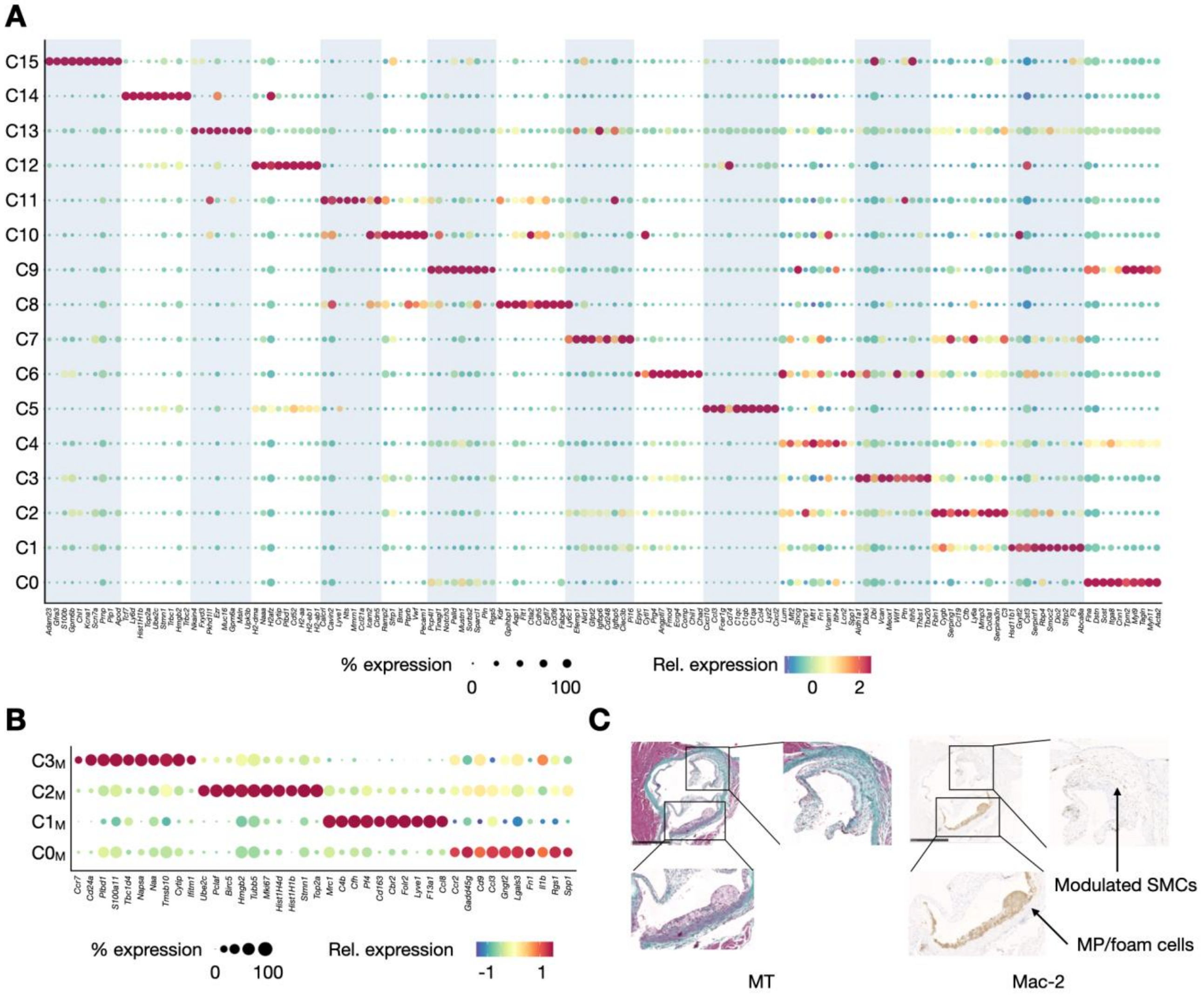


**Figure S14. Single-cell RNA sequencing maps cellular vessel wall and plaque composition [II]. A.** Dot plot showing the top differentially expressed genes for each cluster. **B**. Dot plot showing the top 10 differentially expressed genes for each myeloid cell subcluster. **C**. Representative images of atherosclerotic lesions in aortic valves from treated $WD_{6w}$ mice stained with Masson's Trichrome (MT) and Mac-2 antibody (scale bar = 500 µm). The magnifications [2x] show two types of Mac-2$^{+}$ cells, namely MP/foam cells and modulated SMCs.
